# Reversible, Sexually Dimorphic Sympathetic Aging: Distal Axon Withdrawal Shaped by Prepubertal Gonadal Influences

**DOI:** 10.64898/2026.09.03.748995

**Authors:** Gabriella Rita Pangilinan, Sagar Patil, Francisco Mires, George Ye, Victoria Bogomilova, Lorraine Tilley, Tali Goldkorn, Cassandra J. McGill, Nina Ballou, Xue Li, Unmesh Jadhav, Bérénice A. Benayoun, Yulia Shwartz

## Abstract

Peripheral innervation declines with age, but whether this process is sexually dimorphic and whether it reflects neuronal degeneration or deterioration of nerve–target interactions remain unclear. Using mouse skin as a defined neuroeffector system, we show that sympathetic aging is strongly sexually dimorphic, with greater denervation in aged males. Skin-projecting sympathetic neurons remain intact, indicating that denervation reflects distal axon withdrawal rather than neuronal loss. Nerve–target interactions and the arrector pili muscle niche deteriorate with age, predominantly in males. Aged sympathetic neurons retain NGF responsiveness, and restoring NGF within arrector pili muscles promotes sympathetic reinnervation and hair regeneration in aged males. Genetic and endocrine models show that divergent aging trajectories are associated with gonadal rather than chromosomal sex and are attenuated by early gonadectomy. Together, these findings identify sympathetic aging as a reversible, sexually dimorphic distal axonopathy shaped by the target environment and gonadal influences.

## Introduction

The sympathetic nervous system innervates virtually every organ and is essential for maintaining tissue homeostasis throughout life. Beyond its classical roles in cardiovascular regulation, thermoregulation, metabolism, and digestion, sympathetic nerves dynamically adapt tissue function to environmental and physiological demands. Sympathetic nerves have also emerged as important regulators of tissue regeneration and adult stem cell function, indicating that peripheral nerves contribute directly to tissue maintenance rather than simply controlling organ physiology ^1–4^.

Increasing evidence suggests that the sympathetic nervous system itself undergoes substantial remodeling with age. Age-associated loss or disruption of sympathetic innervation has been reported in multiple organs, including the bone, bone marrow, gastrointestinal tract, heart, and thymus, where it is associated with impaired tissue maintenance and regenerative capacity ^5–10^. Yet the mechanisms underlying this decline remain poorly understood. In particular, whether age-associated denervation reflects degeneration of sympathetic neurons themselves or deterioration of the target tissues that maintain their peripheral axons remains unresolved.

Aging is also profoundly sexually dimorphic. Males and females differ in lifespan, regenerative capacity, susceptibility to age-related disease, and peripheral neuropathy ^11–15^, while sympathetic activity and autonomic regulation exhibit well-established sex differences throughout life ^16–19^. Despite these differences, whether peripheral sympathetic innervation follows distinct aging trajectories in males and females – and what establishes such trajectories – remain unknown.

Peripheral sympathetic innervation depends on continuous communication between neurons and their target tissues. During development, target-derived neurotrophic factors, extracellular matrix components, and specialized stromal cells establish tissue-specific patterns of sympathetic innervation, and these signals continue to support axon maintenance and remodeling throughout adulthood ^20–23^. Among them, nerve growth factor (NGF) is a central regulator of sympathetic neuron survival and target innervation ^24–26^. Peripheral sympathetic axons also retain substantial structural plasticity, undergoing target-dependent sprouting, retraction, and regeneration in response to changes in trophic support and the local tissue environment^27–29^. These observations raise the possibility that age-associated sympathetic decline reflects deterioration of the neuroeffector niche rather than irreversible neuronal loss, but this possibility has not been directly tested.

The skin provides a tractable system to address this question because sympathetic nerves form anatomically defined neuroeffector units with arrector pili muscles (APMs) and hair follicle stem cells (HFSCs). APMs support hair follicle-associated sympathetic fibers, while sympathetic-derived norepinephrine regulates HFSC activation and hair regeneration in response to systemic and environmental cues ^30–33^. This stereotyped organization enables direct visualization and manipulation of neuron–target interactions, providing a tractable model for dissecting mechanisms that may more broadly govern sympathetic aging.

Here, we use the skin as a model to determine how peripheral sympathetic innervation ages, whether this process differs between the sexes, and whether age-associated denervation reflects neuronal degeneration or deterioration of the target niche. We show that sympathetic innervation declines in both sexes but substantially more in males, through distal axon retraction rather than loss of sympathetic neurons. Neuroeffector disorganization precedes overt differences in nerve abundance and is accompanied by sexually dimorphic structural and molecular remodeling of the target niche. Restoring NGF specifically within aged APMs promotes sympathetic reinnervation and is associated with improved hair regeneration, demonstrating that substantial age-associated denervation remains reversible. Finally, in chemical and genetic models, we show that divergent aging trajectories track with gonadal sex rather than sex chromosome complement and implicate early-life gonadal influences in their establishment.

## Results

### Sympathetic innervation declines more severely in the skin of aged males

To ask whether cutaneous sympathetic innervation ages in a sex-dependent manner, we examined tyrosine hydroxylase (TH) positive nerves in thick (100 μm) skin sections from young and old (20 months) male and female C57BL/6JNia mice. Consistent with our previous work, sympathetic axons in young skin formed a dense interconnected network composed of longitudinal axons running alongside hair follicles and transverse branches connecting neighboring follicles (^32^; Fig. 1A). Aging disrupted this network in both sexes, but the extent of degeneration differed markedly. In old females, transverse axons frequently appeared thinner and fragmented while maintaining much of the network architecture. In contrast, old males exhibited extensive loss of transverse branches, resulting in large gaps within the sympathetic network (Fig. 1A).

**Figure 1.**
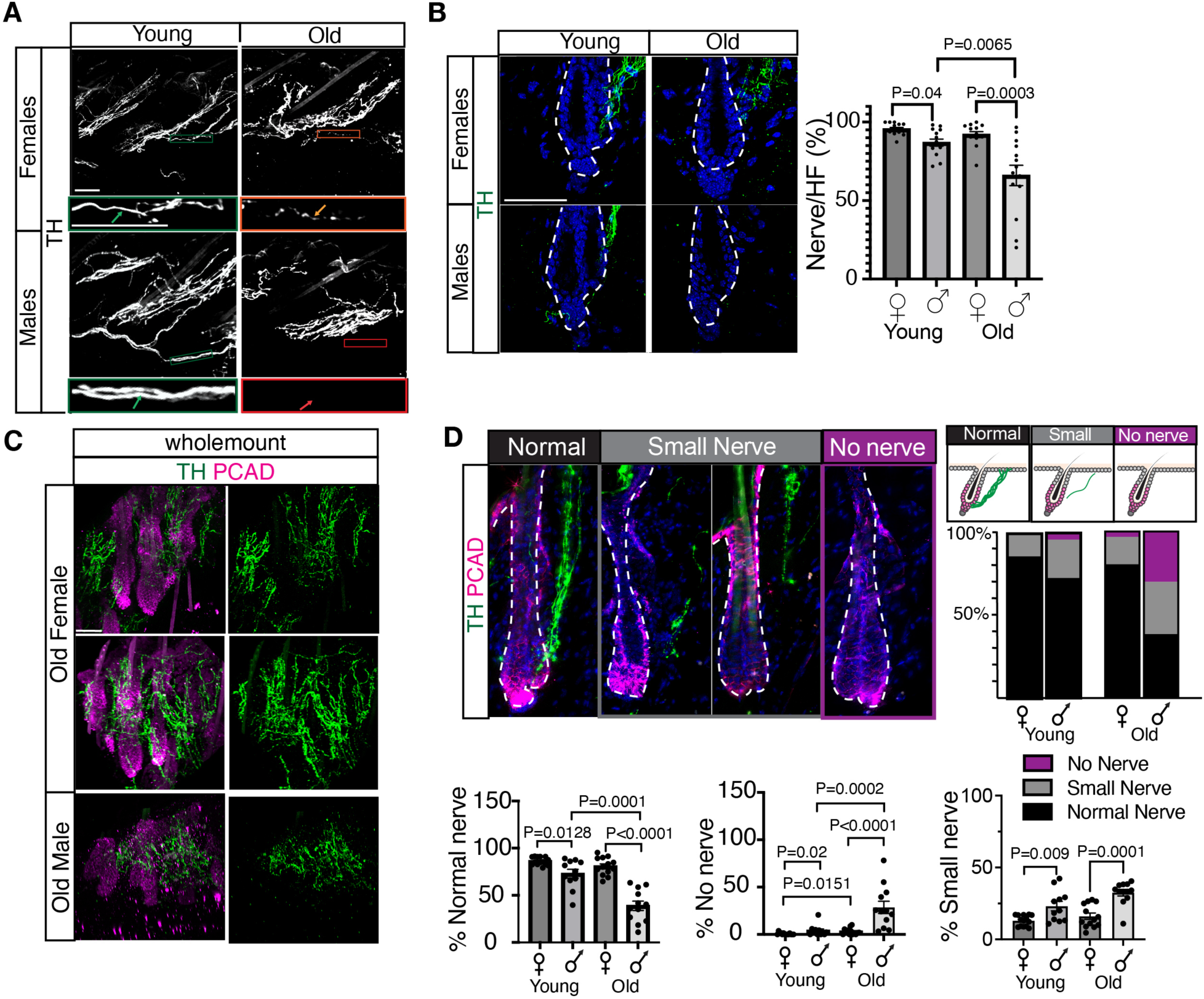
| Aging causes a male-biased decline in sympathetic innervation of hair follicles. (A) Representative images of tyrosine hydroxylase (TH) immunostaining in 100-µm-thick skin sections from young and old female and male mice. Insets show higher-magnification views highlighting intact transverse sympathetic axons in young skin and thinning or fragmented axons in aged skin. Scale bar, 50 µm. (B) Representative TH immunostaining of hair follicle-associated sympathetic nerves and quantification of the percentage of innervated hair follicles (Nerve/HF). (C) Representative whole-mount images of TH-positive sympathetic nerves (green) and P-cadherin (PCAD; magenta) showing hair follicle sympathetic innervation in aged female and aged male skin. Scale bar, 50 µm. (D) Representative TH and P-cadherin (PCAD) immunostaining illustrating the three categories used for quantitative analysis: Normal nerve, Small nerve, and No nerve. Dashed lines outline individual hair follicles. Top right, schematic illustrating the classification criteria. Middle right, distribution of hair follicles classified as Normal nerve, Small nerve, or No nerve in young and old female and male mice. Bottom, quantification of hair follicles exhibiting Normal nerve (left) or No nerve (right). Unless otherwise indicated, each dot represents one mouse (n = 11–14 mice per group), with 30–80 hair follicles analyzed per mouse. Bars represent mean ± s.e.m. Statistical significance was determined using two-sided Mann–Whitney U tests. Exact P values are shown.

Because longitudinal sympathetic axons directly associate with individual hair follicles and regulate hair follicle stem cells (HFSCs), we next quantified hair follicle innervation. Nearly all hair follicles in young males and females were associated with a TH-positive nerve, although young males showed a small but significant reduction compared with females. This difference became substantially more pronounced with age: in old males, approximately 35% of hair follicles completely lacked sympathetic innervation, whereas most hair follicles in old females remained innervated (Fig. 1B). Thus, sympathetic innervation declines with age predominantly in males.

### Surviving sympathetic terminals deteriorate more severely in aged males

Because sympathetic signaling depends on terminal organization and target engagement as well as axon presence, we next asked whether sympathetic nerves that persist with age undergo structural remodeling. Whole-mount imaging of TH together with the hair follicle marker P-cadherin (PCAD) revealed that sympathetic terminals in old males were substantially smaller than those in young animals or age-matched females (Fig. 1C). Three-dimensional reconstruction confirmed a significant reduction in terminal nerve volume in aged males. In addition, many remaining axons were displaced farther from the hair follicle epithelium (Fig. S1A-B).

To quantify these changes, we classified each hair follicle according to the morphology of its associated sympathetic nerve (Fig. 1D). **Normal nerves** consisted of multiple thick terminal axons contacting the follicle, whereas **small nerves** consisted of only one or two thin terminal axons that often failed to contact the follicle. Hair follicles lacking detectable TH-positive axons were classified as **no nerve**. Consistent with the innervation analysis, modest differences in terminal morphology were already evident between young males and females and became substantially greater with age. Approximately 80–90% of hair follicles in young animals displayed normal sympathetic innervation. This proportion remained ∼80% in old females but declined to ∼38% in old males, accompanied by increased frequencies of small and absent nerves (Fig. 1D).

To determine whether this degeneration extended to other cutaneous nerve populations, we examined hair follicle-associated sensory fibers and Merkel cell innervation. TUJ1-positive sensory nerves remained associated with hair follicles in both sexes during aging, and Merkel cell-associated nerves were largely maintained, although occasional denervated Merkel cells were observed in old males (Fig. S1C,D). Thus, age-associated structural deterioration was substantially more pronounced in sympathetic nerves than sensory nerves.

### Aging disrupts sympathetic terminal phenotype and neuroeffector organization

We next asked whether aging alters the phenotype and neuroeffector organization in surviving sympathetic terminals. Hair follicle-associated sympathetic axons remained noradrenergic, as demonstrated by co-expression of tyrosine hydroxylase (TH) and dopamine β-hydroxylase (DBH) (Fig. S2A). We therefore examined the sympathetic co-transmitters neuropeptide Y (NPY) and vasoactive intestinal peptide (VIP), which mark functionally distinct sympathetic populations ^21,34,35^. The proportion of TH-positive axons expressing NPY declined significantly in old males but was largely maintained in females (Fig. 2A). In contrast, VIP expression was unchanged with age or sex (Fig. S2B), indicating selective, sex-dependent changes in sympathetic co-transmitter expression.

**Figure 2.**
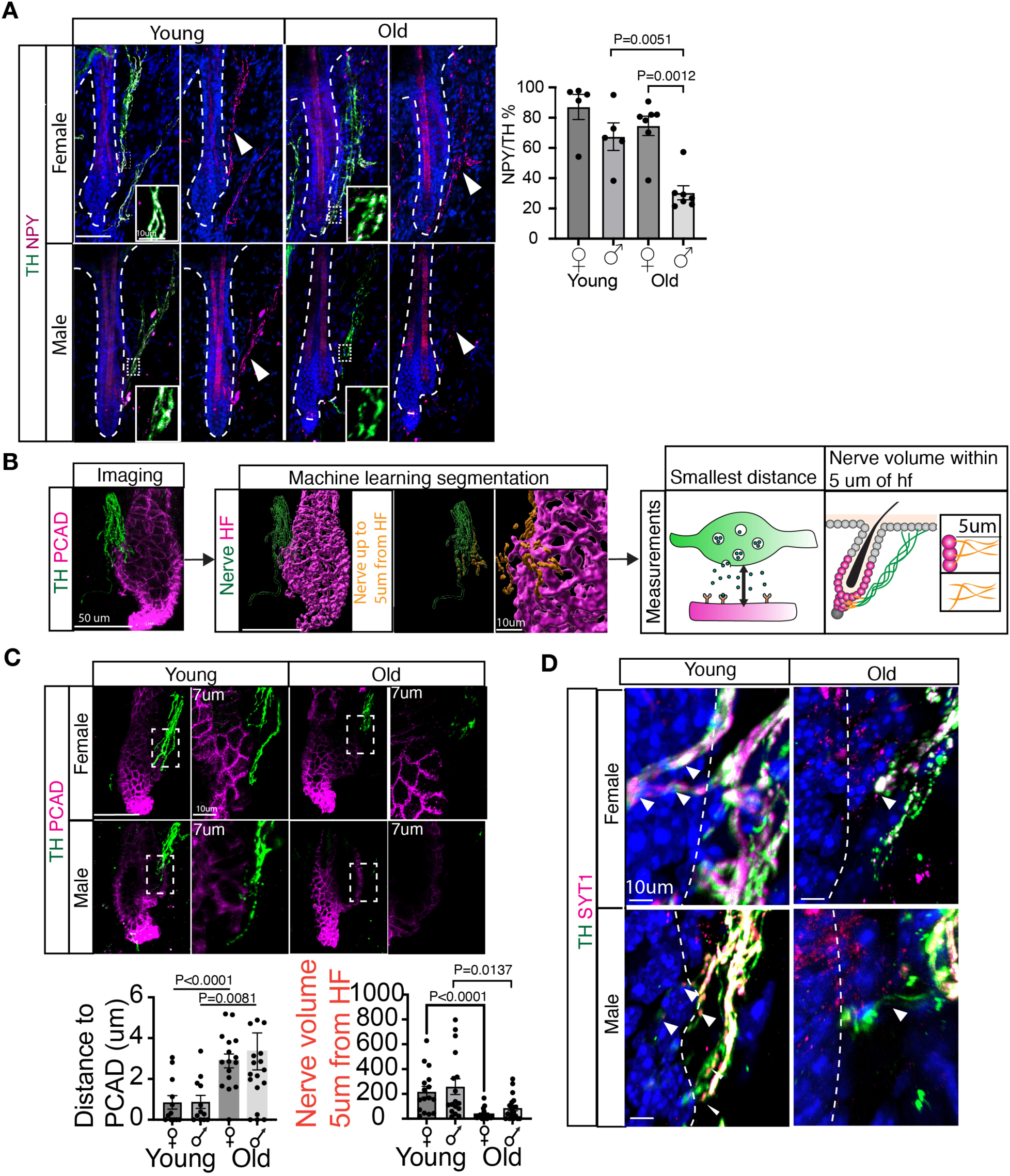
| Aging weakens neuroeffector interactions and presynaptic organization. (A) Representative whole-mount images of tyrosine hydroxylase (TH; green) and neuropeptide Y (NPY; magenta) immunostaining in young and old female and male skin. Arrowheads indicate TH-positive fibers lacking detectable NPY immunoreactivity. Insets show higher-magnification views of representative sympathetic axons. (B) Workflow for three-dimensional image analysis. Whole-mount images of TH-positive sympathetic nerves (green) and P-cadherin (PCAD)-labeled hair follicles (magenta) were segmented using a machine learning-based pipeline to quantify the shortest distance between sympathetic nerves and hair follicles and the volume of sympathetic nerve fibers located within 5 µm of the hair follicle. (C) Representative whole-mount images and higher-magnification views showing sympathetic nerve localization relative to hair follicles in young and old female and male skin. Bottom, quantification of the shortest distance between sympathetic nerves and hair follicles (left) and sympathetic nerve volume within 5 µm of the hair follicle (right). (D) Representative whole-mount images of TH-positive sympathetic nerves (green) and synaptotagmin-1 (SYT1; magenta) immunostaining in young and old female and male skin. Arrowheads indicate representative SYT1-positive presynaptic puncta. Scale bars, 10 µm. For (A), each dot represents one mouse (n = 5–7 mice per group), with 40–150 sympathetic nerve terminals analyzed per mouse. For (C), n = 3 mice per group, with 3–6 images analyzed per mouse; each dot represents one image. Bars represent mean ± s.e.m. Statistical significance was determined using two-sided Mann–Whitney U tests. Exact P values are shown. Scale bars: (A), 50 µm; insets, 10 µm. (B), 50 µm; segmentation inset, 10 µm. (C), 50 µm; higher-magnification images, 10 µm.

Because sympathetic signaling to HFSCs depends on diffusion of norepinephrine and neuropeptides across neuroeffector junctions, we next examined the spatial relationship between sympathetic axons and hair follicles ^36,37^. Three-dimensional reconstructions and machine learning-based segmentation were used to quantify the shortest nerve-to-follicle distance and nerve volume within 5 μm of the follicle (Fig. 2B). Aging increased nerve-to-follicle distance and reduced nerve volume within this perfollicular zone (Fig. 2C), consistent with impaired neuroeffector coupling.

We also examined the presynaptic vesicle protein Synaptotagmin-1 (SYT1) as a measure of presynaptic specialization. In young skin, TH-positive axons displayed abundant SYT1-positive puncta near hair follicles. In aged male skin, SYT1 labeling was markedly diminished along the remaining sympathetic axons (Fig. 2D), suggesting reduced presynaptic specialization.

Collectively, these findings show that surviving sympathetic terminals undergo molecular and organizational changes with age that are more pronounced in males.

### Neuroeffector disorganization precedes overt sympathetic nerve loss

To determine when sympathetic innervation begins to deteriorate and when sex differences first emerge, we analyzed hair follicle-associated sympathetic nerves in male and female mice from 2 to 15 months of age. The percentage of hair follicles with adjacent sympathetic nerves and terminal nerve area did not differ significantly with age or sex through 15 months (Fig. S3A-C). However, 3D reconstruction and machine learning-based segmentation revealed earlier changes in neuroeffector organization. By 15 months, sympathetic nerves in males were positioned significantly farther from hair follicles than those in age-matched females (Fig. S3D), despite comparable nerve number and area. This increased nerve-to-follicle distance was accompanied by reduced sympathetic nerve volume within the 5-μm perfollicular zone (Fig. S3E). Thus, sex-dependent disruption of neuroeffector organization is evident before the pronounced male-biased loss of sympathetic innervation observed at advanced age.

### Axon retraction rather than neuronal loss underlies age-related sympathetic denervation of the skin

Because loss of skin innervation could reflect either neuronal loss or withdrawal of peripheral axons, we asked whether aged ganglia retain skin-projecting sympathetic neurons. Sympathetic neurons innervating the skin reside in the paravertebral sympathetic chain, where individual ganglia also project to other peripheral organs^38^. We therefore used intradermal injection of retrogradely transported AAV expressing GFP or tdTomato ^39^ to selectively label skin-projecting sympathetic neurons within the broader population of TH-positive sympathetic neurons (Fig. 3A). We validated the reproducibility and specificity of this labeling approach across injection conditions (Fig. S4).

**Figure 3.**
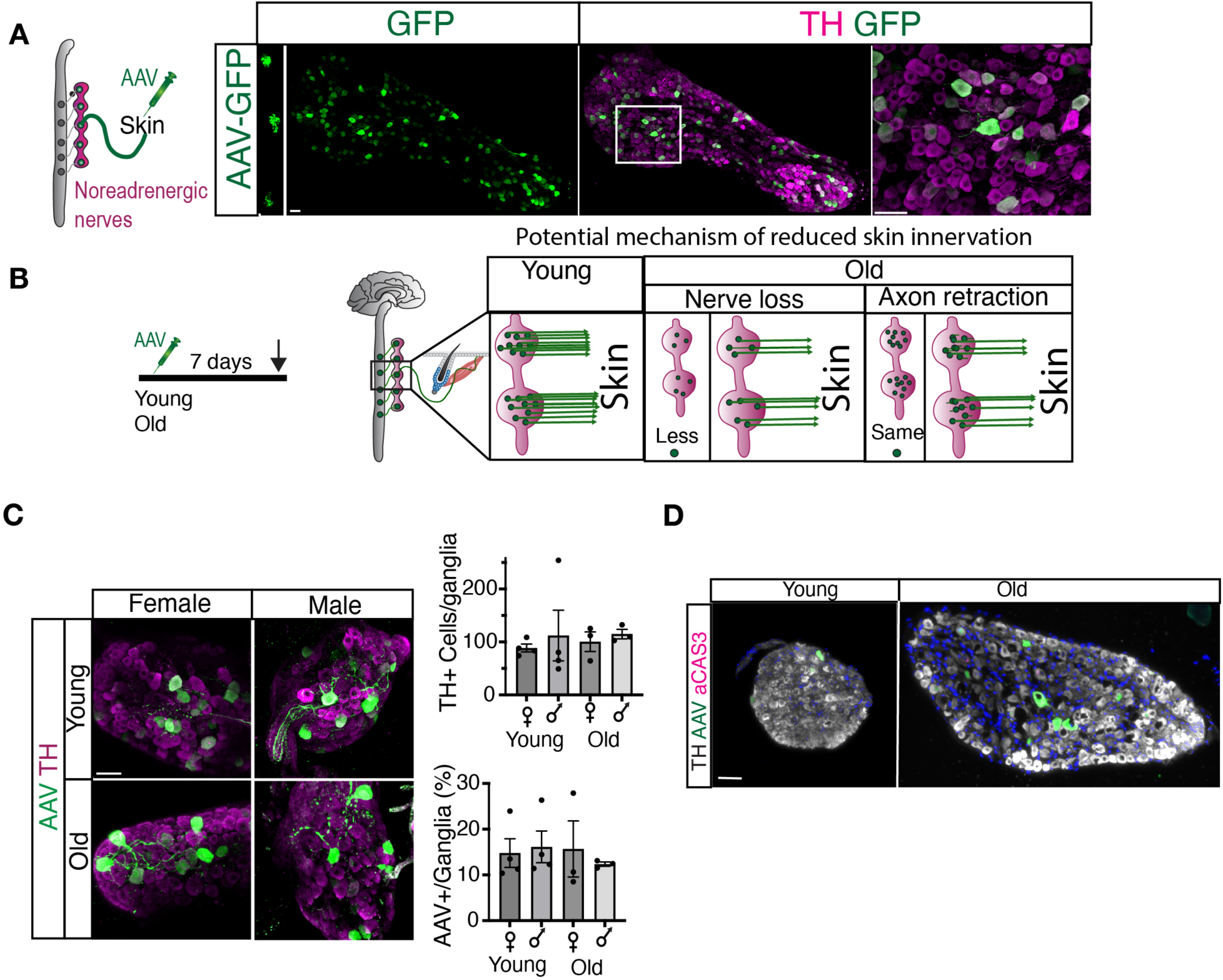
| Skin-projecting sympathetic neurons are preserved during aging. (A) Schematic illustrating retrograde labeling of skin-projecting sympathetic neurons by intradermal injection of AAV-GFP. Representative images of GFP-positive neurons (green) and tyrosine hydroxylase (TH; magenta) immunostaining in thoracic sympathetic ganglia. Right, higher-magnification view of the boxed region. (B) Experimental design for retrograde labeling of skin-projecting sympathetic neurons in young and old mice. Right, schematic illustrating two potential mechanisms underlying reduced skin sympathetic innervation during aging: neuronal loss or distal axon retraction. (C) Representative images of GFP-positive neurons (green) and TH immunostaining (magenta) in thoracic sympathetic ganglia from aged female and aged male mice. Right, quantification of the total number of TH-positive neurons per ganglion (top) and the percentage of GFP-positive skin-projecting sympathetic neurons (bottom). Each dot represents a different mouse. (D) Representative images of activated caspase-3 (aCAS3; magenta), TH (white), and GFP (green) immunostaining in thoracic sympathetic ganglia from young and old mice Scale bar, 50 µm for all images. For (C), 3-4 mice were analyzed per group, with 1–5 ganglia analyzed per mouse. Bars represent mean ± s.e.m. Statistical significance was determined using two-sided Mann–Whitney U tests. Exact P values are shown.

AAV-retro was injected into the dorsal skin of young and old mice after the aging phenotype was fully established (20 months; Fig. 3B). We first asked whether the total sympathetic neuron population changes with age. Whole-mount TH staining of the anterior thoracic ganglia revealed no significant differences in sympathetic neuron numbers between young and old mice of either sex (Fig. 3C), consistent with previous observations in human and animal sympathetic ganglia ^7,40^. We then quantified retrogradely labeled skin-projecting neurons following a single intradermal AAV-retro injection, which consistently labels neurons within the T1-T5 sympathetic ganglia.

Furthermore, the percentage of GFP-positive neurons was comparable between young and old animals of both sexes (Fig. 3C), indicating that skin-projecting sympathetic neurons are maintained during aging.

Consistent with neuronal preservation, activated caspase-3 (aCAS3) staining in males showed no evidence of increased apoptosis in aged sympathetic ganglia (Fig. 3D). Together, these findings indicate that the age-related loss of cutaneous sympathetic innervation reflects distal axon retraction rather than loss of sympathetic neuronal cell bodies.

### Arrector pili muscle degeneration accompanies but does not fully explain sympathetic denervation

Because APMs provide structural support for hair follicle-associated sympathetic nerves ^32^, we asked whether APM aging could contribute to sympathetic denervation. Previous work identified age-associated changes in APM structure in females ^41^, but whether this process differs between sexes is unknown. Using integrin α8 (ITGA8) staining, we found that APMs in young animals typically formed continuous fibers connecting the hair follicle to the epidermis, whereas aging was associated with structural abnormalities, including APM loss, detachment from the epidermis or hair follicle, and breaks within the muscle (Fig. S5A).

Quantification at 2, 15 and 20 months revealed progressive APM remodeling in both sexes, with substantially greater deterioration in males (Fig. S5B,C). By 15 months, males exhibited fewer normal APMs and increased frequencies of APM loss and detachment, with these differences becoming more pronounced by 20 months. A weighted APM degeneration index that accounted for the severity of these structural abnormalities (Methods) similarly increased with age in both sexes but was significantly higher in males at 15 and 20 months (Fig. S5B).

To determine whether APM degeneration alone could explain sympathetic denervation, we examined APM morphology in denervated hair follicles from old males. Only 30.8% of denervated follicles had completely lost the APM, while an additional 14% showed detachment from the hair follicle (Fig. S5D). Most denervated follicles retained an APM attached to the HFSC compartment but detached from the epidermis. Thus, structural degeneration of the APM accompanies age-related sympathetic denervation but is insufficient to explain nerve loss, suggesting that additional changes within the target niche contribute to denervation.

### NGF declines more prominently in aged male skin

Because structural changes in the niche did not fully explain denervation, we next asked whether sympathetic decline was associated with reduced trophic support. Nerve growth factor (NGF), a key regulator of sympathetic neuron survival and maintenance^26^, declined with age in both sexes but was reduced more prominently in aged males (Fig. 4A), consistent with the greater sympathetic denervation observed in male skin. To determine whether local NGF abundance predicted nerve integrity, we assessed sympathetic nerve morphology within NGF-positive APMs. In aged females, NGF-positive APMs were predominantly associated with normal sympathetic nerve morphology, whereas in aged males they were frequently associated with small or absent nerves (Fig. 4B). Thus, reduced NGF accompanies the greater sympathetic decline in aged males, but NGF abundance alone does not predict maintenance of sympathetic innervation.

**Figure 4.**
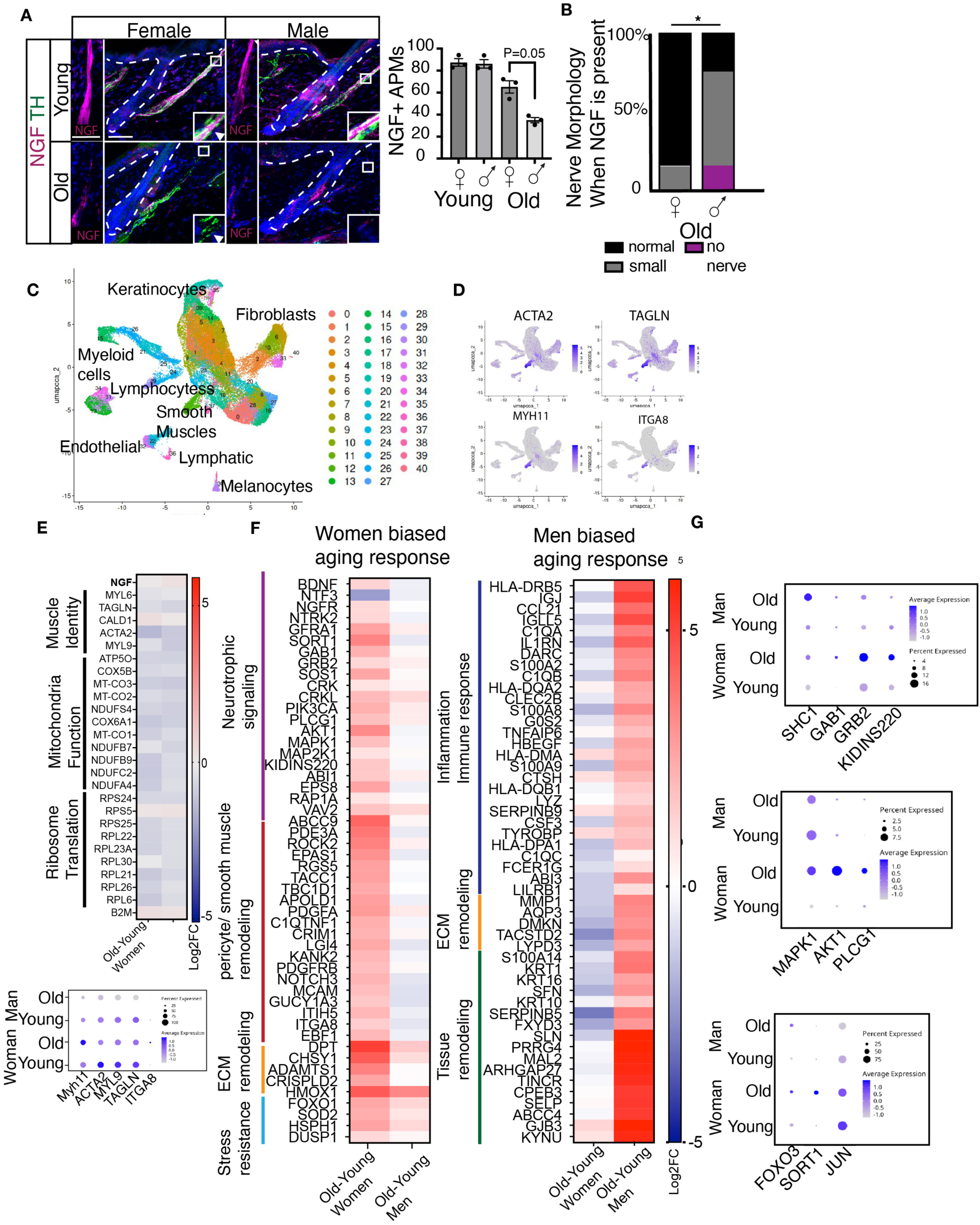
| Aging remodels neurotrophic support and smooth muscle transcriptional programs in a sex-dependent manner. (A) Representative images of nerve growth factor (NGF; magenta) and tyrosine hydroxylase (TH; green) immunostaining in young and old female and male skin. Dashed lines outline individual hair follicles. Insets show higher-magnification views of NGF-positive arrector pili muscles (APMs). Right, quantification of the percentage of NGF-positive APMs (left). (B) Distribution of sympathetic nerve morphology associated with NGF-positive APMs. (C) Uniform manifold approximation and projection (UMAP) of the integrated human skin single-cell RNA-seq dataset showing the major skin cell populations. (D) Feature plots showing expression of ACTA2, TAGLN, MYH11, and ITGA8, identifying the smooth muscle cell cluster. (E) Heatmap showing representative genes exhibiting conserved age-associated transcriptional changes in smooth muscle cells from women and men, including smooth muscle identity, mitochondrial oxidative phosphorylation, and ribosomal pathways. Bottom, dot plot representative expression of MYH11, TAGLN, ACTA2, MYL9 and ITGA8. Dot size represents the percentage of expressing cells and color indicates scaled average expression. (F) Heatmaps showing representative genes exhibiting women-biased (left) and men-biased (right) age-associated transcriptional responses in smooth muscle cells. (G) Dot plots showing representative components of the neurotrophin signaling pathway exhibiting relatively higher expression in smooth muscle cells from aged women than aged men. Dot size represents the percentage of expressing cells and color indicates scaled average expression. For (A), each dot represents one mouse (n = 3 mice per group), with 70-100 hair follicles analyzed per mouse. For (B), each bar represents the mean distribution across three mice per group. Bars represent mean ± s.e.m. Statistical significance was determined using two-sided Mann–Whitney U tests. Exact P values are shown. Scale bars: (A), 50 µm

### Human skin smooth muscle undergoes sex-dependent transcriptional remodeling with age

To determine whether sex-dependent remodeling of the smooth muscle compartment also occurs during human aging, we integrated published single-cell RNA-seq datasets from young, middle-aged, and aged women and men ^42–44^ (Fig. *S6A,B*). Unsupervised clustering identified major skin cell populations, including a smooth muscle cluster defined by *ACTA2, MYH11, TAGLN*, and *ITGA*8 expression (Fig. 4C,D; Fig. S6C). Because these datasets do not distinguish arrector pili from vascular smooth muscle, we interpret this population as a general skin smooth muscle compartment.

We compared age-associated transcriptional changes in smooth muscle cells from women and men and identified both shared and sex-biased changes (Methods; Table1,2). Based on differences in the magnitude of age-associated fold changes, 7,075 genes were classified as showing women-biased age-associated changes, 4,744 as showing men-biased age-associated changes, and 3,384 as showing shared age-associated regulation. Canonical smooth muscle identity genes (*ACTA2, MYH11, MYL9, TAGLN, and ITGA8*) showed modest age-associated reductions that were broadly similar between sexes. Mitochondrial oxidative phosphorylation and ribosomal genes likewise declined with age in both women and men (Fig. 4E). *NGF* transcript levels also changed little with age, suggesting that the age-associated decline in NGF protein is not primarily reflected at the transcript level.

Sex-biased changes were evident in pathways associated with neurotrophic support, inflammation and tissue remodeling (Fig. 4F,G) ^45–50^. Age-associated changes in women favored multiple components of neurotrophin signaling, including receptors and co-receptors (*NGFR, NTRK2, GFRA1, SORT1*), intracellular signaling mediators (*PIK3CA, AKT1, MAPK1, PLCG1*) and associated cytoskeletal regulators. Genes associated with pericyte identity and extracellular matrix regulation were also relatively enriched in aged women.

By comparison, age-associated changes in men favored antigen-presentation and complement genes, inflammatory mediators, and genes associated with epithelial and tissue remodeling (Fig. 4F). Thus, human skin smooth muscle undergoes sexually dimorphic transcriptional remodeling with age, characterized by relative preservation of neurotrophin-associated programs in women and greater inflammatory and tissue-remodeling programs in men.

### APM-derived NGF promotes sympathetic reinnervation in aged male skin

To determine whether reduced neurotrophic support contributes to age-related sympathetic axon retraction, we first asked whether skin-projecting sympathetic neurons retain the capacity to respond to NGF during aging. Retrograde labeling of skin-projecting neurons with TrkA immunostaining showed that sympathetic neurons expressed the high-affinity NGF receptor TrkA across age and sex (Fig. 5A). Moreover, phosphorylated TrkA (p-TrkA) levels were comparable across groups (Fig. 5B), indicating that aged sympathetic neurons remain competent to respond to NGF signaling.

**Figure 5.**
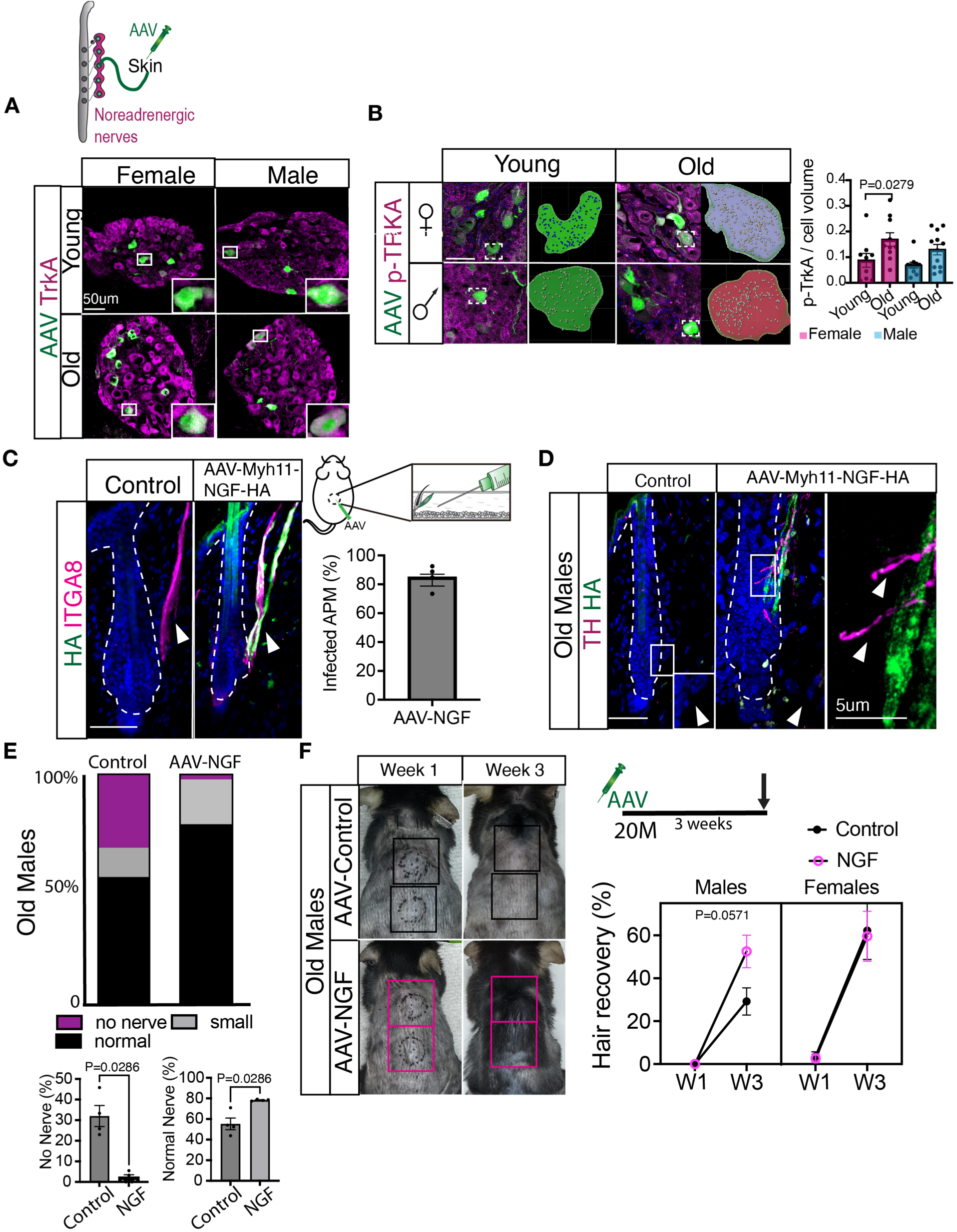
| NGF overexpression restores sympathetic innervation and is associated with enhanced hair regeneration in aged skin. (A) Schematic illustrating retrograde labeling of skin-projecting sympathetic neurons by intradermal injection of AAV-GFP. Representative images of GFP-positive neurons (green) and TrkA immunostaining (magenta) in thoracic sympathetic ganglia from young and old female and male mice. Insets show higher-magnification views of representative GFP-positive neurons. (B) Representative images of phosphorylated TrkA (p-TrkA; magenta) and GFP-positive neurons (green) in thoracic sympathetic ganglia from young and old female and male mice. Right, quantification of p-TrkA immunoreactivity normalized to neuronal volume. (C) Experimental strategy for arrector pili muscle (APM)-specific NGF overexpression using AAV-Myh11-NGF-HA. Representative whole-mount images of HA (green) and integrin α8 (ITGA8; magenta) immunostaining showing selective transduction of APMs. Right, quantification of infected APMs. (D) Representative images of tyrosine hydroxylase (TH; magenta) and HA (green) immunostaining in aged male skin following control or AAV-Myh11-NGF-HA treatment. Insets show higher-magnification views of TH-positive sympathetic nerve terminals associated with HA-expressing APMs. (E) Distribution of hair follicles classified as Normal nerve, Small nerve, or No nerve following AAV-Myh11-NGF-HA treatment. Bottom, quantification of hair follicles exhibiting No nerve (left) and Normal nerve (right). (F) Experimental design for NGF overexpression in aged mice. Representative images showing hair regeneration in control– and AAV-Myh11-NGF-HA-treated aged mice at weeks 1 and 3. Right, quantification of hair recovery in female and male mice. For (B), n = 3 mice per group. Three to four images were analyzed per mouse, with 10–22 GFP-positive skin-projecting neurons analyzed per mouse. Each dot represents the average quantification from one image. For (C) and (F), n = 4 mice per group. Bars represent mean ± s.e.m. Statistical significance was determined using two-sided Mann–Whitney U tests. Exact P values are shown. Scale bars: (A–D), 50 µm; (D) inset, 5 µm.

We next asked whether restoring local NGF could promote reinnervation. As a proof of principle, we broadly overexpressed NGF in aged skin using AAV8-CAG-NGF-GFP, which primarily targeted dermal fibroblasts with occasional expression in APMs. NGF overexpression increased both sympathetic (TH) and sensory (TUJ1) innervation, with newly formed axons frequently localized adjacent to NGF-expressing cells (Fig. S7A). Thus, aged cutaneous nerves retain the capacity to respond to locally increased NGF.

Because broad NGF expression also increased sensory innervation, we generated an APM-specific AAV in which the smooth muscle *Myh11* promoter drives expression of HA-tagged NGF (AAV-Myh11-NGF-HA). This construct selectively targeted approximately 80% of APMs within the injected region (Fig. 5C). Importantly, APM-specific NGF expression did not alter neighboring hair follicle-associated sensory innervation (Fig. S7B), allowing us to assess the effects of restoring NGF specifically within the APM niche.

In old males, APM-specific NGF expression promoted close association of sympathetic axons with NGF-expressing APMs (Fig. 5D), increased the proportion of hair follicles with normal sympathetic innervation, and reduced the proportion of denervated follicles. No major effect on sympathetic innervation was observed in aged females (Fig. 5E; Fig. S7C). Reinnervation was accompanied by greater hair recovery in aged males, whereas no additional effect was observed in aged females (Fig. 5F). Thus, restoring NGF expression specifically within the aged APM niche promotes sympathetic reinnervation and is associated with improved hair regeneration, demonstrating that age-associated denervation retains substantial regenerative potential.

### Gonadal sex and early-life gonadal hormone exposure shape sexually dimorphic sympathetic aging

Sex differences can arise from circulating sex hormones or sex chromosome complement. Because estrogen can modulate sympathetic function and innervation ^51,52^, we first asked whether preservation of sympathetic nerves in aged females depends on continued ovarian hormone exposure during adulthood. We induced ovarian failure using vinylcyclohexene diepoxide (VCD) ^53^, which depletes ovarian follicles while preserving ovarian tissue, and examined sympathetic innervation at 6 and 20 months (Fig. 6A and Fig S8A). Despite long-term depletion of ovarian follicles, neither the percentage of innervated hair follicles nor terminal nerve area differed between VCD– and vehicle-treated animals at either age (Fig. 6A, Fig. S8A), indicating that post-pubertal ovarian function is not required to maintain sympathetic skin innervation during aging.

**Figure 6.**
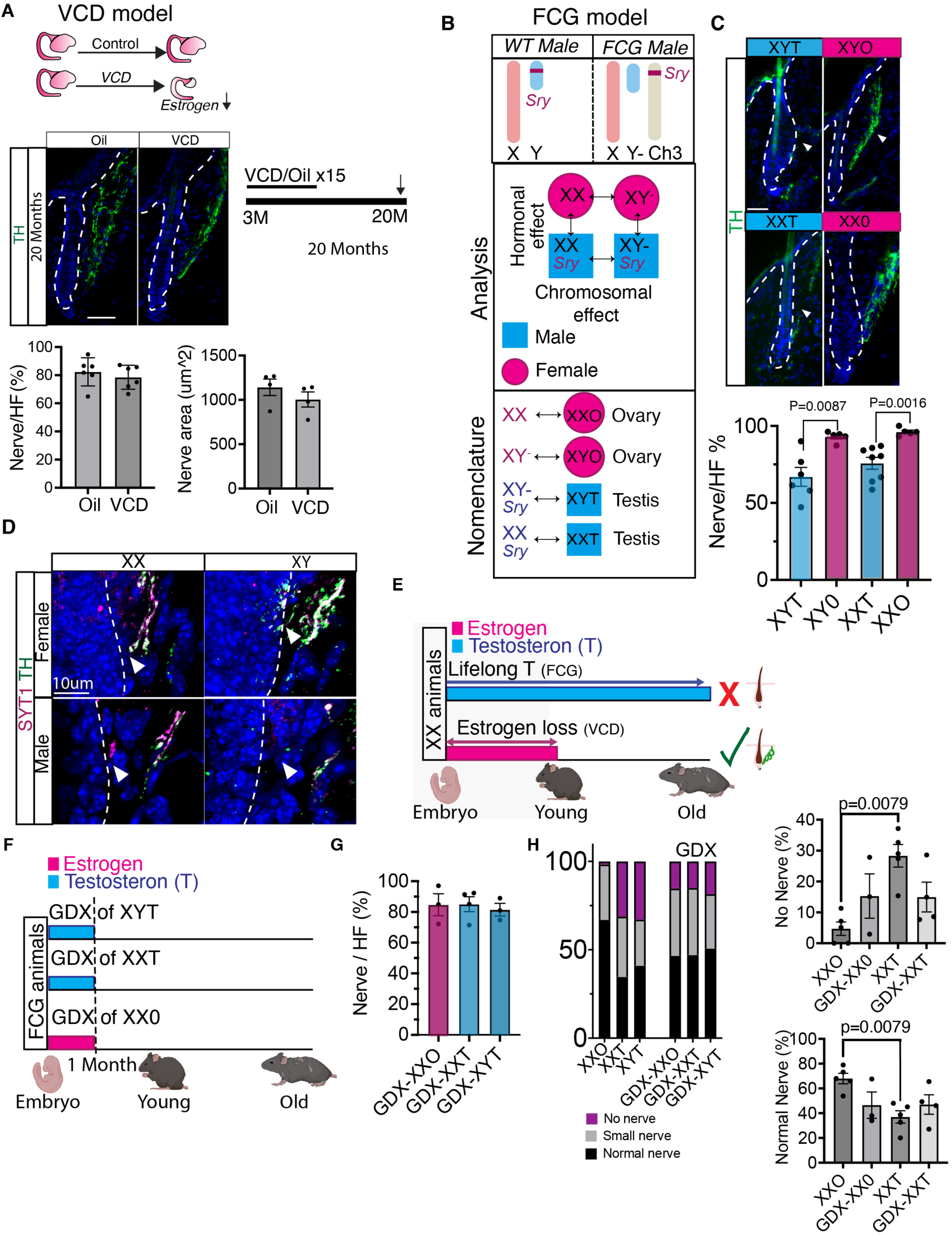
| Gonadal sex determines susceptibility to age-associated sympathetic denervation. (A) Experimental design of the 4-vinylcyclohexene diepoxide (VCD) model of ovarian failure. Representative whole-mount images of tyrosine hydroxylase (TH; green) immunostaining in vehicle– and VCD-treated skin from 20-month-old mice. Right, quantification of the percentage of innervated hair follicles (Nerve/HF; top) and sympathetic nerve area (bottom) in control and VCD-treated mice at 20 months. (B) Schematic of the Four Core Genotypes (FCG) mouse model illustrating the separation of gonadal sex and sex chromosome complement. (C) Representative images of TH immunostaining in aged Four Core Genotypes mice. Right, quantification of the percentage of innervated hair follicles (Nerve/HF). (D) Representative whole-mount images of tyrosine hydroxylase (TH; green) and synaptotagmin-1 (SYT1; magenta) immunostaining in aged Four Core Genotypes mice. Arrowheads indicate representative SYT1-positive presynaptic puncta. Insets show higher-magnification views. Scale bar, 10 µm. (E) Summary model illustrating the contributions of gonadal hormones and sex chromosomes to age-associated sympathetic denervation. (F) Experimental design: XYT, XXT, and XXO mice underwent gonadectomy (GDX) at 1 month and were analyzed at 20-22 months. Testis-bearing animals underwent orchiectomy, and ovary-bearing mice underwent ovariectomy. (G) Quantification of the percentage of innervated hair follicles (Nerve/HF) in GDX mice. (H) Quantification of nerve morphology distribution in 20-22 months old FCG mice with and without GDX. For (A), n = 4–6 mice per group. For the percentage of innervated hair follicles, 20–60 hair follicles were analyzed per mouse. For sympathetic nerve area, 10 images were analyzed per mouse. Each dot represents one mouse. For (C), n = 5–8 mice per group, with 25–60 hair follicles analyzed per mouse. For (G,H), 3–5 mice per group were analyzed, with 20–120 hair follicles analyzed per mouse. Each dot represents one mouse. Bars represent mean ± s.e.m. Statistical significance was determined using two-sided Mann–Whitney U tests. Exact P values are shown. Scale bars: (A, C), 50 µm; (D), 10 µm.

We next used the Four Core Genotypes (FCG) model^54–56^, which uncouples XX/XY chromosome complement from gonadal sex, to determine which factor predicts the sexually dimorphic aging phenotype (Fig. 6B). In aged gonad-intact FCG mice, animals with testes exhibited significantly lower sympathetic innervation than animals with ovaries, whereas animals with the same gonadal sex showed comparable innervation regardless of sex chromosome complement (Fig. 6C,E,H). Thus, the sexually dimorphic decline in sympathetic innervation tracks with gonadal sex rather than XX/XY chromosome complement.

The gonadal sex-dependent phenotype extended beyond nerve abundance: aged mice with testes displayed reduced presynaptic SYT1 labeling, diminished sympathetic innervation of blood vessels, and sensory nerve alterations comparable to those observed in aged wild-type males (Fig. 6D; Fig. S8C-D). To determine whether early-life gonadal exposure contributes to divergent aging trajectories, we performed gonadectomy (GDX) in FCG mice at 1 month of age and examined sympathetic innervation at 20 months (Fig. 6F). Early GDX substantially attenuated differences among FCG groups. The percentage of innervated hair follicles as well as nerve morphology was broadly comparable across XXO, XXT and XYT groups (Fig. 6G,H).

Notably, aged GDX animals exhibited an intermediate distribution of sympathetic nerve morphologies rather than recapitulating either intact aged females or males. Together with the VCD experiment, these findings distinguish effects of early-life gonadal exposure from continued ovarian function in adulthood. Depletion of ovarian function beginning after puberty did not alter the female aging trajectory, whereas removal of the gonads at 1 month substantially attenuated the later sex difference. Thus, early gonadal hormone exposure contributes to establishing sexually divergent trajectories of sympathetic aging.

## Discussion

The sympathetic nervous system is increasingly recognized as a critical regulator of tissue homeostasis throughout adulthood, yet how sympathetic innervation itself ages remains poorly understood. Here, we identify sympathetic aging as a sexually dimorphic process characterized by distal axon retraction rather than neuronal loss. Using retrograde tracing, we directly demonstrate preservation of the tissue-projecting neurons whose peripheral axons are lost with age, while identifying deterioration of the neuroeffector niche as an important contributor to this process. Sex-dependent niche remodeling is evident in both mouse and human skin, and restoring NGF specifically within the APM niche promotes sympathetic reinnervation and is associated with improved hair regeneration in aged males. Finally, genetic and endocrine models reveal that divergent aging trajectories are associated with gonadal rather than chromosomal sex and implicate gonadal influences before puberty in establishing later susceptibility to sympathetic degeneration (Fig. 7).

**Figure 7.**
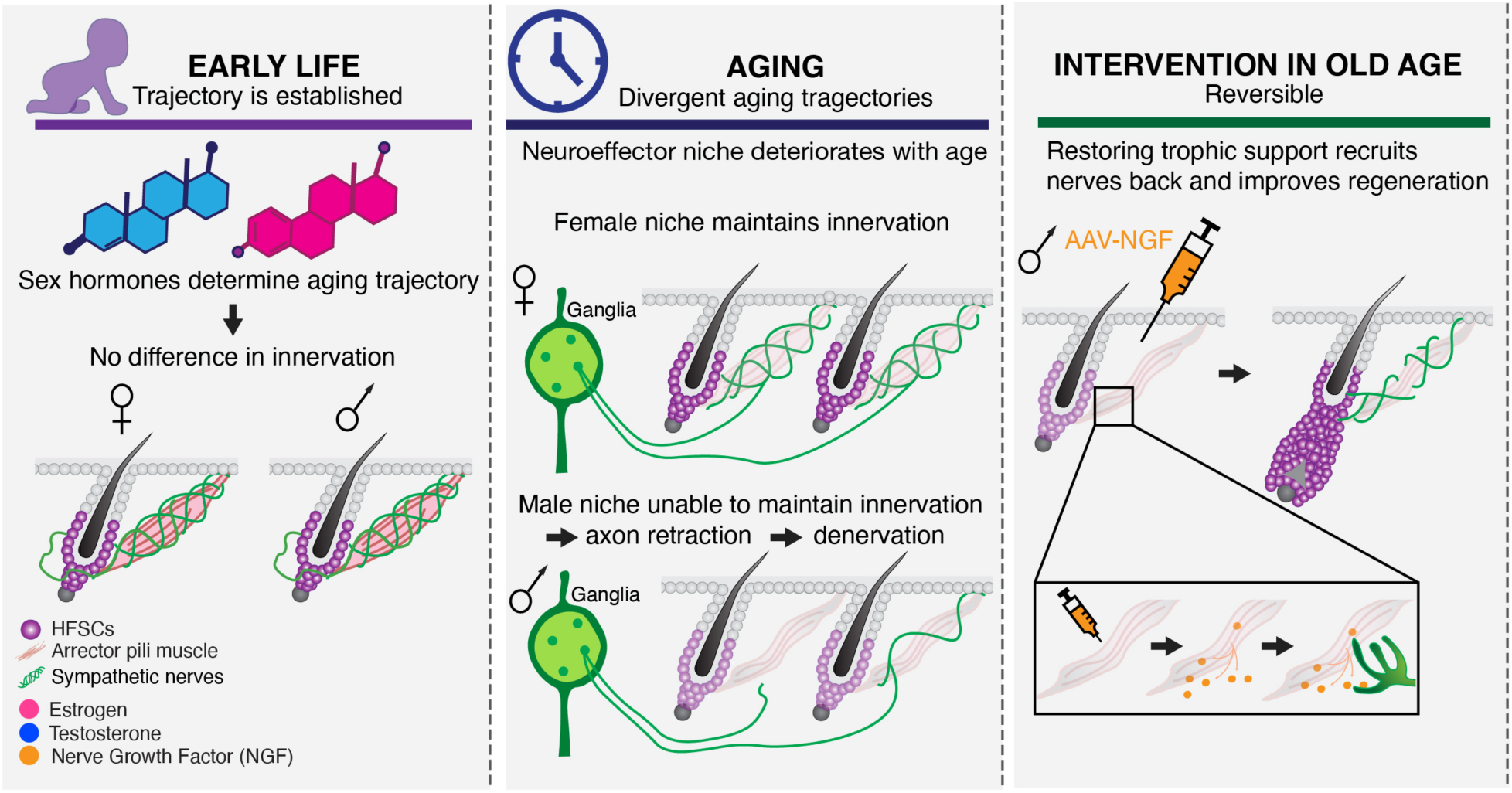
| Model of sexually dimorphic sympathetic aging and its reversibility. Gonadal hormone exposure early in life establishes divergent trajectories of sympathetic aging despite similar initial cutaneous innervation. With age, deterioration of the neuroeffector niche is associated with progressive distal sympathetic axon retraction that is substantially more pronounced in males, while sympathetic neuronal cell bodies remain preserved. Restoring local trophic support through AAV-mediated NGF expression in the arrector pili muscle of aged males promotes sympathetic reinnervation and is associated with enhanced hair regeneration. Together, these findings suggest that early-life gonadal influences shape later susceptibility to sympathetic aging, while preservation of responsive sympathetic neurons allows substantial peripheral denervation to remain reversible.

Age-associated remodeling of sympathetic innervation has been described in the bone marrow, heart, gastrointestinal tract and thymus, where it is associated with impaired tissue maintenance and physiological adaptability ^6–10,57^. Yet whether sympathetic aging differs between males and females has received little attention despite well-established sex differences in sympathetic physiology ^58–60^. Pathological studies of human autonomic ganglia have suggested that sympathetic aging may involve distal axon degeneration despite preservation of neuronal cell bodies ^40^. By retrogradely tracing the sympathetic neurons that specifically innervate the affected skin, we directly demonstrate that these tissue-projecting neurons remain preserved despite substantial loss of their peripheral terminals. Moreover, they retain TrkA signaling competence and the capacity to reinnervate their targets. Thus, sympathetic aging can represent a reversible distal axonopathy rather than irreversible degeneration of the neuron itself.

This distinction shifts attention toward the target tissue as an active determinant of peripheral nerve aging. In the heart, changes in the local tissue environment regulate sympathetic denervation and reinnervation following injury ^61,62^. In the aging heart, endothelial senescence increases the repulsive axon-guidance cue SEMA3A, whereas reducing senescent cells restores innervation and improves autonomic and cardiac function ^8^. In the bone marrow, sympathetic denervation contributes to hematopoietic stem cell niche aging, while restoration of β3-adrenergic signaling improves aged stem cell function ^5^. Thymic aging similarly involves sympathetic axon degeneration together with loss of innervation-supporting stromal populations^9^. Together with our findings, these studies support a model in which sympathetic aging can emerge from progressive disruption of reciprocal interactions between peripheral axons and their target tissues rather than acting exclusively through neuron-intrinsic degeneration.

The hair follicle provides a particularly informative context for this model because HFSC aging is strongly influenced by the surrounding microenvironment ^63–68^. Aged HFSCs largely retain their molecular identity and can recover substantial regenerative capacity in a younger or more supportive environment ^41,69^. Our findings extend this niche-centered view to the neural component of the HFSC environment. Sympathetic nerves, APMs and HFSCs form an integrated neuroeffector unit, and aging progressively disrupts its organization and communication. Thus, aging of the stem cell, its stromal niche and its innervation may represent interconnected features of deterioration within a common regenerative unit rather than independent processes.

A key dimension of this model is that the neuroeffector niche itself ages differently between the sexes. APMs undergo progressive structural remodeling in both sexes, but abnormalities become substantially more pronounced in males. Yet many denervated follicles retain an APM, indicating that structural deterioration alone cannot explain nerve loss. Instead, structural changes occur alongside molecular remodeling, including a more pronounced decline in NGF protein in aged male skin. Importantly, sexually dimorphic aging is also evident in human skin smooth muscle. Although the available datasets do not distinguish APMs from vascular smooth muscle cells and are therefore correlative, aged female smooth muscle cells relatively preserve neurotrophin-associated programs, whereas aged male cells preferentially exhibit inflammatory and tissue-remodeling programs. The parallel between mouse and human data suggests that sex-dependent remodeling of the smooth muscle compartment during aging is not restricted to the mouse model.

NGF is likely one component of a broader landscape governing sympathetic maintenance, and endogenous NGF abundance alone does not predict nerve integrity. NGF-positive APMs in aged males can still be associated with small or absent nerves, and human smooth muscle cells remodel multiple components of neurotrophin signaling despite relatively little change in NGF transcript abundance. Sympathetic aging may therefore reflect a changing balance among trophic, structural, inflammatory and axon-guidance signals rather than loss of a single factor.

Nevertheless, our rescue experiments demonstrate that manipulating one component of this environment can be sufficient to produce substantial reinnervation. Unlike approaches that restore downstream adrenergic signaling in the bone marrow and thymus or broadly modify the aged environment in the heart ^5,8,9^, APM-specific NGF expression manipulates a defined trophic signal within a defined peripheral target cell. The resulting reinnervation demonstrates that even after substantial distal withdrawal, aged sympathetic axons retain the capacity to respond to an improved target environment.

The accompanying improvement in hair regeneration should nevertheless be interpreted with some caution. NGF and its receptors are expressed within hair follicles, and neurotrophin signaling has been implicated in follicular epithelial proliferation and hair-cycle regulation ^70,71^, in addition to its well-established effects on cutaneous innervation ^25^. NGF could therefore influence regeneration through neural, epithelial or coordinated mechanisms. However, the sex specificity of the response parallels the neural phenotype: aged males, which exhibit substantially greater sympathetic denervation, show both reinnervation and improved regeneration following NGF expression, whereas aged females largely retain sympathetic innervation and derive little additional benefit. This association supports a contribution of restored innervation, although direct and nerve-mediated effects of NGF remain to be distinguished.

The capacity of aged sympathetic axons to reinnervate their targets is consistent with the substantial plasticity of peripheral sympathetic circuits including their capacity for neurotrophin-dependent axon regeneration following injury ^72,73^. In inflammatory skin disease, sympathetic fibers undergo context-dependent changes in density, norepinephrine availability and neuronal activity ^74–76^. Our findings extend this plasticity to aging, showing that sympathetic axons remain responsive to changes in their target environment even after substantial distal withdrawal. Local signals governing nerve–target interactions may therefore represent an accessible point for restoring peripheral neural function in aged tissues.

The pronounced sexual dimorphism of sympathetic aging raises a second question: when are these divergent trajectories established? The VCD, Four Core Genotypes and gonadectomy experiments argue against a simple model in which continued adult ovarian hormones protect female innervation. Depletion of ovarian function beginning after puberty did not accelerate denervation, whereas aged FCG animals with testes exhibited greater sympathetic degeneration than those with ovaries irrespective of sex chromosome complement, implicating gonadal rather than chromosomal sex. Strikingly, gonadectomy at 1 month, before puberty and long before overt sympathetic aging, substantially attenuated the later sex difference, with animals converging toward an intermediate phenotype rather than acquiring an intact male– or female-like state. Together, these findings suggest that gonadal influences before puberty contribute to establishing the later trajectory of sympathetic aging.

These experiments do not distinguish whether ovarian hormones confer long-term protection, testicular hormones promote later degeneration, or both contribute. Moreover, because gonadectomy was performed at 1 month rather than before all gonadal hormone exposure, the intermediate phenotype may reflect organizational effects established before gonadectomy that are not fully reversible. Thus, an aging phenotype that becomes pronounced late in life may be shaped, at least in part, by endocrine signals encountered before puberty rather than simply by cumulative exposure to adult sex hormones. This possibility extends emerging niche-centered models of sex differences in regenerative aging, in which endothelial, stromal, inflammatory, vascular and neural components collectively shape stem cell function ^77,78^, by suggesting that gonadal signals may establish sex-specific properties of the regenerative environment long before overt aging phenotypes appear.

Such developmental programming is consistent with broader links between early-life maturation and late-life aging. Pubertal timing in humans is associated with biological aging, frailty and age-related disease ^79–82^, while genetic manipulation of *vgll3*, a conserved regulator of pubertal timing, accelerates male growth and sexual maturation but increases age-related disease and mortality in killifish ^83^. Although these studies do not directly address sympathetic innervation, they support a broader principle in which developmental programs establish persistent tissue states with consequences that emerge later in life. Future studies using gonadectomy at different developmental stages, temporally controlled hormone replacement and tissue-specific manipulation of androgen and estrogen signaling will be required to define the relevant window and determine whether these effects are mediated through sympathetic neurons, the target niche or systemic endocrine pathways.

An important unresolved question is how aging of the different components of the neuroeffector unit influences one another. HFSC function depends on both stromal and sympathetic inputs, but our experiments do not establish whether sympathetic withdrawal contributes to HFSC aging, whether aging of HFSCs or APMs destabilizes innervation, or whether these processes evolve reciprocally. The ability of aged HFSCs to recover regenerative capacity in a younger environment ^41,69^, together with our finding that modifying the aged target niche can restore neural input and improve regeneration, argues against age-related dysfunction arising solely from irreversible deterioration of any single compartment. Instead, regenerative aging may emerge from progressive disruption of communication among stem cells, stromal cells and nerves.

Collectively, our findings support a model in which sympathetic aging reflects progressive disruption of nerve–target interactions rather than irreversible neuronal degeneration, and in which this trajectory differs substantially between the sexes. Direct retrograde tracing demonstrates that tissue-projecting neurons remain preserved despite peripheral axon loss, while sexually dimorphic remodeling of the target niche in mouse and human identifies sex as an important determinant of the environment encountered by aging axons. The attenuation of these differences by prepubertal gonadectomy further suggests that susceptibility to late-life sympathetic degeneration is shaped, at least in part, before overt aging begins. Together with the ability to restore neural input by modifying the aged niche, these findings suggest that peripheral nerve aging emerges from interactions among developmental history, the aging target environment and persistent neuronal plasticity (Fig. 7). More broadly, the ability of aged HFSCs to recover in a younger environment and our ability to restore neural input by modifying the aged niche support a model in which regenerative aging is a property of the integrated neuron–target–stem cell unit rather than of any single cellular compartment.

## Author Contributions

G.R.P. conceived and designed experiments, performed experiments, analyzed and interpreted data, and contributed to writing the manuscript. S.P., F.M. and V.B. performed experiments and analyzed data. G.Y. analyzed the published human single-cell RNA-sequencing datasets. T.G. contributed to analysis of existing experimental datasets. C.J.M. and N.B. performed experiments and contributed to the study, with N.B. also providing Four Core Genotype mice. X.L. supervised N.B. and provided scientific guidance. U.J. supervised G.Y. and contributed to discussions of the single-cell analyses. B.A.B. provided the aged mouse cohorts used in the initial studies as well as VCD-treated cohorts and contributed to scientific discussions and interpretation. Y.S. conceived and supervised the study, designed experiments, interpreted data and wrote the manuscript. All authors discussed the results and commented on the manuscript.

## Acknowledgements

We thank the members of the Shwartz laboratory and our collaborators for helpful discussions and technical assistance. We thank Seth Ruffins, Arkadi Shwartz, and the Optical Imaging Facility at USC Stem Cell and the Translation Imaging Center for assistance with microscopy and image analysis. We thank the USC and Cedars-Sinai animal facilities for animal husbandry.

We thank Kristen Mehalko for performing VCD injections. We also thank the investigators who generated and made publicly available the human skin single-cell RNA-sequencing datasets analyzed in this study.

## Materials and Methods

### Mice

All animal procedures were approved by the Institutional Animal Care and Use Committee (IACUC) at the University of Southern California (USC) and Cedars-Sinai and were performed in accordance with institutional and NIH guidelines. Studies were conducted under USC IACUC protocols 21489 (Shwartz lab), 20770, 20804, 21004, 21454, and 21214 (Benayoun lab), and IACUC009544 (Li lab) as applicable.

C57BL/6J mice were used unless otherwise indicated. Unmanipulated 20-month-old wild-type mice used for aging comparisons were obtained from the NIA aged rodent colony; unmanipulated 2-month– and 15-month-old mice were obtained from Jax. All manipulated animals, including VCD experiments and AAV-based experiments, were obtained from The Jackson Laboratory.

Four Core Genotype (FCG) mice were originally obtained from Arthur P. Arnold (UCLA) and aged at USC or Cedars-Sinai. All FCG mice examined contained the previously described 3.2-Mb X-to-Y chromosome translocation ^84^.

Mice were housed under specific-pathogen-free conditions in individually ventilated cages with food and water available ad libitum and, unless otherwise indicated, a 12-h light/12-h dark cycle. Both sexes were included. For sympathetic innervation analyses, mice 2–5 months of age were operationally grouped as young because cutaneous sympathetic innervation did not differ detectably across this age range (Fig. S3A–C); this grouping does not imply equivalent developmental or endocrine status. Exact ages are provided in the figure legends. Unless otherwise stated, mice ≥20 months were considered old.

#### VCD model of ovarian failure

To induce accelerated ovarian follicle depletion, mice received intraperitoneal injections of 4-vinylcyclohexene diepoxide (VCD; Sigma-Aldrich, cat. no. 94956) at 160 mg kg−1 in safflower oil once daily for 15 consecutive days, starting at 3 months of age. Vehicle controls received matched volumes of safflower oil alone. Body weight and general health were monitored throughout treatment. Mice were euthanized and analyzed at the indicated time points after treatment.

#### Gonadectomy

Four Core Genotypes (FCG) mice underwent gonadectomy between postnatal day 21 and 35 (P21–P35). Mice with ovaries underwent bilateral ovariectomy, and mice with testes underwent bilateral orchiectomy. Animals were anesthetized with isoflurane or ketamine/xylazine, and depth of anesthesia was confirmed before surgery. Buprenorphine-XR was administered for perioperative analgesia. For ovariectomy, the ovaries were accessed through dorsal or bilateral flank incisions, the ovarian vessels and uterine horns were ligated, and both ovaries were removed. For orchiectomy, the testes were accessed through a lower abdominal incision and removed bilaterally following cauterization of the associated vasculature. Incisions were closed with absorbable sutures and wound clips or nylon sutures. Animals were maintained on a heating pad during recovery and monitored postoperatively. Wound clips or sutures were removed 7–10 days after surgery. Gonadectomized animals were subsequently aged to 20-22 months for analysis of cutaneous innervation.

#### Viral vectors and intradermal administration

Readily available AAV constructs were purchased from Addgene or Boston Children’s Viral Core. NGF overexpression viruses were generated and packaged by Welgen. For broad NGF overexpression, AAV8-CAG-GFP served as control and AAV8-CAG-NGF-T2A-GFP was used for overexpression. For arrector pili muscle-specific NGF delivery, AAV-PHP.S-Myh11-NGF-HA was used. For retrograde labeling of skin-projecting sympathetic neurons, AAV-retro-CAG-GFP was used. Viral stocks were diluted in sterile saline to 1 × 10^12 genome copies (gc) ml−1 and injected intradermally into the dorsal skin at a final dose of 4–5 × 10^10 gc per injection. One to three intradermal injections per mouse were used. Viruses were co-injected with fluorescent beads to determine the injection location. Skin was collected 7 days after AAV-retro-CAG-GFP injection and 2 months after AAV8-CAG-GFP, AAV8-CAG-NGF-T2A-GFP or AAV PHP.S-Myh11-NGF-HA injection.

#### Tissue collection and immunofluorescence

Dorsal skin was fixed in 4% paraformaldehyde for 15 min at room temperature, washed in PBS, cryoprotected in 30% sucrose overnight at 4°C, embedded in OCT and sectioned at 50-100 µm. Sympathetic ganglia were processed similarly and sectioned at 10 µm. Sections were blocked in PBS containing 5% donkey serum, 1% bovine serum albumin, 2% cold-water fish gelatin and 0.3% Triton X-100 for 1–2 h at room temperature, and incubated with primary antibodies overnight at 4 °C followed by fluorophore-conjugated secondary antibodies for 3–4 h at room temperature or overnight at 4 °C. For 100 µm, thick sections, incubation with both primary and secondary antibodies was prolonged to 2 days at 4 °C. For NGF staining, additional antigen retrieval using 10 mM sodium citrate, pH 6.0 was used before blocking and incubation with primary antibody, as well as Tyramide Signal Amplification (TSA) plus Fluorescein staining after the secondary antibody incubation step.

For skin whole-mount staining, dorsal skin was fixed in 4% paraformaldehyde, dehydrated through a methanol series, bleached in 5% hydrogen peroxide, rehydrated, and blocked at 37°C for 4 h. Samples were immunolabeled with 2-day antibody incubations at 4 °C and cleared in benzyl alcohol/benzyl benzoate (BAB) before imaging. The full list of primary antibodies and dilutions is provided in the Key Resources Table.

For wholemount ganglia staining: Sympathetic ganglia were collected and fixed in 4% paraformaldehyde for 15 min at room temperature, washed in PBS and dehydrated through a graded methanol series (20%, 40%, 60%, 80% and 100% methanol; 30 min each at room temperature). Samples were subsequently incubated once more in 100% methanol at 4 °C and bleached overnight at 4 °C in 5% hydrogen peroxide in methanol. Ganglia were rehydrated through a graded methanol series (80%, 60%, 40%, 20% and PBS containing 0.3% Triton X-100; 30 min each at room temperature). Samples were permeabilized overnight at 37 °C with shaking in permeabilization buffer containing 0.3% Triton X-100, 2.3% (w/v) glycine and 20% (v/v) DMSO in PBS. Samples were then blocked for 4 h at 37 °C with shaking in PBS containing 5% donkey serum, 1% bovine serum albumin, 2% cold-water fish gelatin and 0.3% Triton X-100. Primary antibodies were incubated for 2 d at 4 °C with shaking, followed by fluorophore-conjugated secondary antibodies for 2 d at 37 °C with shaking. Samples were washed between antibody incubations using PTwH buffer containing 0.3% Tween-20 and 10 µg ml⁻¹ heparin in PBS. The full list of primary antibodies and dilutions is provided in the Key Resources Table.

#### Microscopy and image analysis

Images were acquired using Leica Stellaris, Leica SP8, Zeiss LSM800 confocal microscopes, or a Leica Thunder widefield microscope. Z-stacks were collected and processed as maximum-intensity projections or single optical sections, as indicated in the figure legends. Acquisition settings were kept constant within each experiment.

<u>Hair follicle-associated sympathetic nerves</u> were quantified as the percentage of hair follicles with an adjacent TH-positive nerve or as sympathetic nerve area, depending on the experiment. Hair follicles were classified as having a normal, small, or absent sympathetic nerve according to predefined morphological criteria illustrated in Fig. 1d. For <u>sympathetic nerve area</u> quantifications: Sympathetic nerve area was quantified in Fiji (ImageJ). TH-stained nerves were imaged at ×40 magnification using a widefield microscope with extended depth-of-focus (Full Focus) processing. Individual sympathetic nerves were manually cropped from surrounding tissue prior to automated thresholding. Images were converted to 8-bit grayscale, and segmented using the IJ IsoData automatic threshold. Binary masks were generated, and the area of each TH-positive nerve was measured following image calibration.

<u>Three-dimensional analysis of sympathetic nerve morphology and neuroeffector organization</u> was performed on TH– and P-cadherin (PCAD)-stained skin sections imaged using a Leica Stellaris confocal microscope with an ×87 objective. Confocal image stacks were segmented in Imaris using machine learning-based classification to generate volumetric reconstructions of sympathetic nerves and hair follicles. Identical segmentation parameters were applied to all experimental groups within each experiment. This segmentation pipeline was used to quantify sympathetic nerve volume (Supplementary Fig. S1). For neuroeffector analyses (Fig. 2), a distance transformation was applied to the PCAD-defined hair follicle surface to generate a 5– µm perfollicular zone, and sympathetic nerve volume contained within this region was quantified. The shortest three-dimensional distance between the sympathetic nerve surface and the hair follicle was also measured. Together, nerve volume within the 5-µm perfollicular zone and the shortest nerve-to-hair follicle distance were used as complementary surrogates of local neuroeffector coupling.

<u>p-TrkA quantification</u>. Phosphorylated TrkA (p-TrkA) was quantified in skin-projecting sympathetic neurons following retrograde labeling with AAV-retro-CAG-GFP. Confocal z-stacks were acquired using a Leica Stellaris confocal microscope with an ×87 objective. GFP-positive neuronal cell bodies were segmented in Imaris to generate three-dimensional cell volumes. p-TrkA immunoreactivity was detected using the Imaris Spots function, and the number of p-TrkA-positive puncta was normalized to neuronal volume to obtain p-TrkA spot density. Three animals were analyzed per group, with at least three images acquired per animal and 13–30 GFP-positive neurons analyzed per animal. Each data point represents the mean p-TrkA spot density per image.

<u>Arrector pili muscle (APM) morphology</u> was evaluated in 100-µm skin sections stained for integrin α8 (ITGA8) or smooth muscle α-actin (SMA). Individual hair follicles were classified into one of six predefined categories based on APM morphology: normal, no APM, disconnected from the epidermis, disconnected from the hair follicle, disconnected from both the epidermis and hair follicle, or break in the middle. Classification was performed manually using predefined morphological criteria. To quantify the overall severity of APM degeneration, a weighted degeneration index was calculated from the distribution of these phenotypes. Complete APM loss was assigned the highest score (3), detachment from the hair follicle or simultaneous detachment from both the hair follicle and epidermis were assigned an intermediate score (2), and detachment from the epidermis or a break in the middle were assigned a lower score (1). The weighted degeneration index was calculated as the average severity score across all analyzed hair follicles.

<u>NGF-positive APM quantification:</u> The prevalence of NGF-positive arrector pili muscles (APMs) was quantified in 50-µm skin sections immunostained for smooth muscle α-actin (SMA), tyrosine hydroxylase (TH), and nerve growth factor (NGF). Individual hair follicles were scored based on the presence or absence of an APM, a sympathetic nerve, and NGF expression within the APM. NGF-positive APMs were further classified according to the morphology of their associated sympathetic nerve as normal, small, or absent, using the predefined criteria described above. Three animals were analyzed per group, with 70–100 hair follicles analyzed per animal. For NGF-positive APM quantification, each data point represents the percentage of NGF-positive APMs calculated for an individual animal. For nerve morphology, the distribution of normal, small, and absent sympathetic nerves was calculated for each animal and is presented as the mean across three animals.

Hair regeneration was quantified from standardized dorsal photographs by measuring the percentage of recovered hair at the AAV injection site using Fiji.

Whole-mount ganglia were imaged using a Leica Thunder widefield microscope, and TH-positive neurons and retrogradely labeled GFP-positive neurons were manually quantified in Fiji (ImageJ).

#### Analysis of published human single-cell RNA-seq datasets

Previously published human skin single-cell RNA-sequencing datasets from ^42–44^ were analyzed in R (v4.6.1) using Seurat (v5.5.0). Individual datasets were processed independently by log-normalization, identification of highly variable genes, scaling of gene expression, and principal component analysis (PCA).

Samples within each dataset were integrated using Seurat’s canonical correlation analysis (CCA) workflow, followed by construction of shared nearest-neighbor graphs, graph-based clustering, and Uniform Manifold Approximation and Projection (UMAP) for visualization. Cell populations were annotated using established lineage marker genes. Sex was confirmed using XIST expression together with Y chromosome-associated genes (EIF1AY, DDX3Y, KDM5D, USP9Y, TMSB4Y, NLGN4Y, ZFY and RPS4Y1).

The integrated dataset comprised 56,038 cells, including 5,454 young male, 10,003 aged male, 10,425 young female, 9,846 middle-aged female, and 20,310 aged female cells. Smooth muscle cells were identified as cluster 12 based on expression of ACTA2, MYH11, TAGLN, and ITGA8 and contained 1,515 cells, including 554 young male, 282 aged male, 172 young female, 106 middle-aged female, and 401 aged female cells. Because the analyses focused on age-associated changes, downstream differential expression analyses were restricted to young and aged samples.

Differential gene expression between aged and young smooth muscle cells was performed separately for women and men using Seurat’s Wilcoxon rank-sum test. Gene lists from the two comparisons were matched by gene symbol and classified according to sex differences in age-associated log2 fold change. Genes were classified as women-biased when the absolute age-associated log2 fold change in women exceeded that in men by ≥ 0.5 (|women log2FC| − |men log2FC| ≥ 0.5), men-biased when the converse was true (|men log2FC| − |women log2FC| ≥ 0.5), and shared when both sexes changed in the same direction, the absolute difference in log2 fold change between sexes was ≤ 0.5, and the mean absolute log2 fold change was ≥ 0.5. These criteria identified 7,075 women-biased genes, 4,744 men-biased genes, and 3,384 genes exhibiting conserved age-associated regulation.

Because this study focused on mechanisms regulating neuron–target communication, downstream analyses emphasized genes involved in neurotrophin signaling, smooth muscle identity, extracellular matrix remodeling, and inflammatory pathways. Representative genes were selected based on effect size, biological relevance, and their established roles in neurotrophin signaling or smooth muscle biology.

Data sets used: HRA000395; GSE150672; GSE130973

#### Statistics and reproducibility

Statistical analyses were performed in GraphPad Prism and R. The experimental unit was the mouse unless otherwise indicated. For p-TrkA quantification, image-level means were used for statistical analysis, with the mouse as the biological replicate. For image-based quantifications, multiple images or structures from the same mouse were treated as technical replicates, and the number of biological replicates, images and structures analyzed for each figure is reported in the figure legends. Data are presented as mean ± s.e.m. unless otherwise indicated. Exact P values, sample sizes and statistical tests are provided in the figure legends. Two-sided Mann–Whitney U tests or two-sided Student’s t-tests were used, as appropriate, for the comparisons shown in the figures. WT Animals were randomly assigned to experimental groups when applicable. Image acquisition and quantification were performed blinded to genotype and treatment whenever possible.

| Reagent or Resource | Source | Identifier |
| --- | --- | --- |
| Antibodies |  |  |
| Rabbit anti Tyrosine hydroxylase (1:1000) | Millipore | Cat#ab152 |
| Sheep anti Tyrosine hydroxylase (1:150) | Millipore | Cat#ab1542 |
| Chicken anti Tyrosine hydroxylase (1:100) | Millipore | Cat#ab9702 |
| Guinea pig anti dopamine β-hydroxylase (1:250) | Synaptic Systems | Cat# 496 004 |
| Goat anti Integrin alpha 8 (1:200) | R&D | Cat#AF4076 |
| Rabbit anti Smooth Muscle Actin (1:400) | Abcam | Cat#ab5694 |
| Rabbit anti Tuj1 (1:1000) | Millipore | Cat# T2200 |
| Rabbit anti Synaptotagmin $\frac{1}{2}$ (1:500) | Synaptic Systems | Cat#105003 |
| Rabbit anti HA (1:200) | Cell Signaling Technology | Cat# 3724S |
| Rabbit anti GFP (1:500) | Abcam | Cat#ab290 |
| Rabbit anti Vasoactive intestinal polypeptide (1:250) | Cell Signaling Technologies | Cat# 63269T |
| Rabbit anti Neuropeptide Y (1:250) | Cell Signaling Technology | Cat#11976T |
| Rabbit anti TrkA (1:250) | Sigma | 06-574 |
| Phospho-TrkA (Tyr785)/TrkB (Tyr816) (C67C8) Rabbit mAb #4168 1:250 | Cell Signaling | 4168S |
| Anti- NGF, beta subunit, Purified, (Polyclonal) (sheep IgG) 1:100 | CEDARLANE | CLMCNET-031 |
| Peroxidase AffiniPure® F(ab')<sub>2</sub> Fragment Donkey Anti-Sheep IgG (H+L) | Jackson ImmunoResearch | 713-036-147 |
| TSA Plus Fluorescein Kit | Akoya Biosciences, Quanterix | nel741001kt |
| **Chemicals:** |  |  |
| PBS pH 7.4 (10X) | Gibco | 70011044 |
| Saline (0.9% NaCl) | Sigma | S8776-6X20ML |
| Ethanol | Avantik |  |
| Methanol | Avantik | RS4039 |
| Hydrogen peroxide | Sigma | 1085971000 |
| Donkey Serum | Sigma | D9663-10ML |
| Bovine Serum Albumin | Sigma | A7030-10g |
| Gelatin from cold water fish skin | VWR | 97062-618 |
| Triton X-100 | VWR | 100504-970 |
| 16% Paraformaldehyde | Electron Microscopy Sciences | 15710 |
| Glycine | Sigma | G7126-100G |
| Sucrose | Sigma | S9378-5KG |
| Tween-20 | Sigma | P1379-100ML |
| DMSO | Sigma | D8418-50ML |
| Heparin | Sigma | H3149-10KU |
| VCD | Sigma-Aldrich | 94956 |
| Safflower oil | Sigma | S8281-500ML |
| **Viruses** |  |  |
| AAVrg-CAG-GFP | BCH |  |
| AAVrg-CAG-tdTOMATO | Addgene | Addgene viral prep # 59462-AAVrg |
| AAV-PHP.S-MYH11-NGF-HA | This paper |  |
| AAV8-CAG-NGF-T2A-GFP | <sup>85</sup> |  |

## Supplement

**Supplementary Figure S1.**
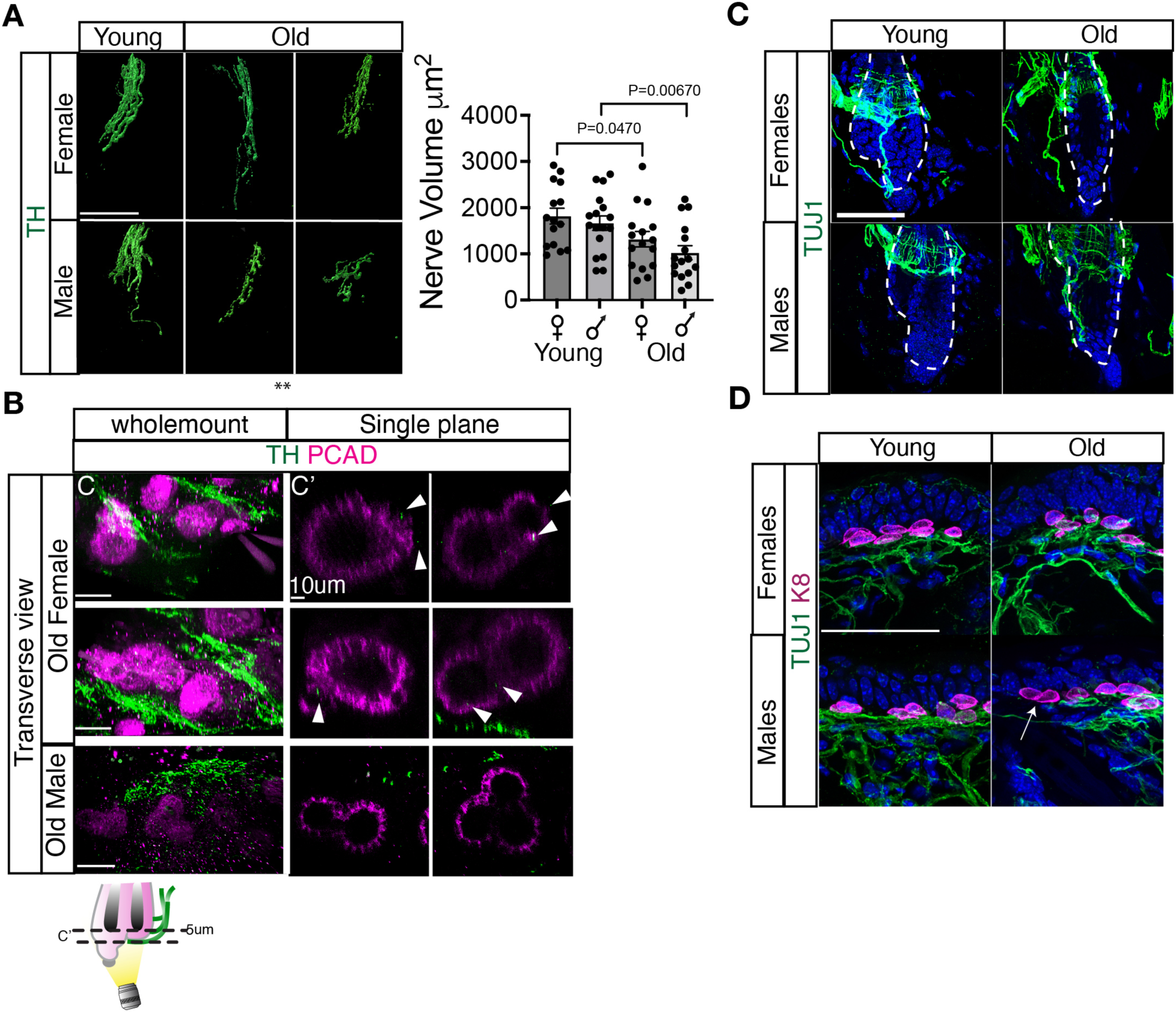
| Aging remodels sympathetic nerve morphology. (A) Three-dimensional reconstructions of tyrosine hydroxylase (TH)-positive sympathetic nerve terminals associated with hair follicles in young and old female and male mice. Right, quantification of sympathetic nerve terminal volume. (B) Representative whole-mount images of TH-positive sympathetic nerves (green) and P-cadherin (PCAD; magenta) in aged female and aged male skin. Left, whole-mount view. Right, single optical sections through the hair follicle illustrating sympathetic nerve terminal morphology. Arrowheads indicate representative sympathetic nerve terminals. Schematic illustrates the imaging plane. (C) Representative images of TUJ1 immunostaining showing hair follicle-associated sensory nerves in young and old female and male skin. (D) Representative images of TUJ1 (green) and keratin 8 (K8; magenta) immunostaining showing Merkel cell-associated sensory innervation in young and old female and male skin. For (A), n = 3 mice per group, with 4–6 images analyzed per mouse. Each dot represents one image. Bars represent mean ± s.e.m. Statistical significance was determined using two-sided Student’s t-tests. Exact P values are shown. Scale bars: (A–D), 50 µm; (B) single optical sections, 10 µm.

**Supplementary Figure S2.**
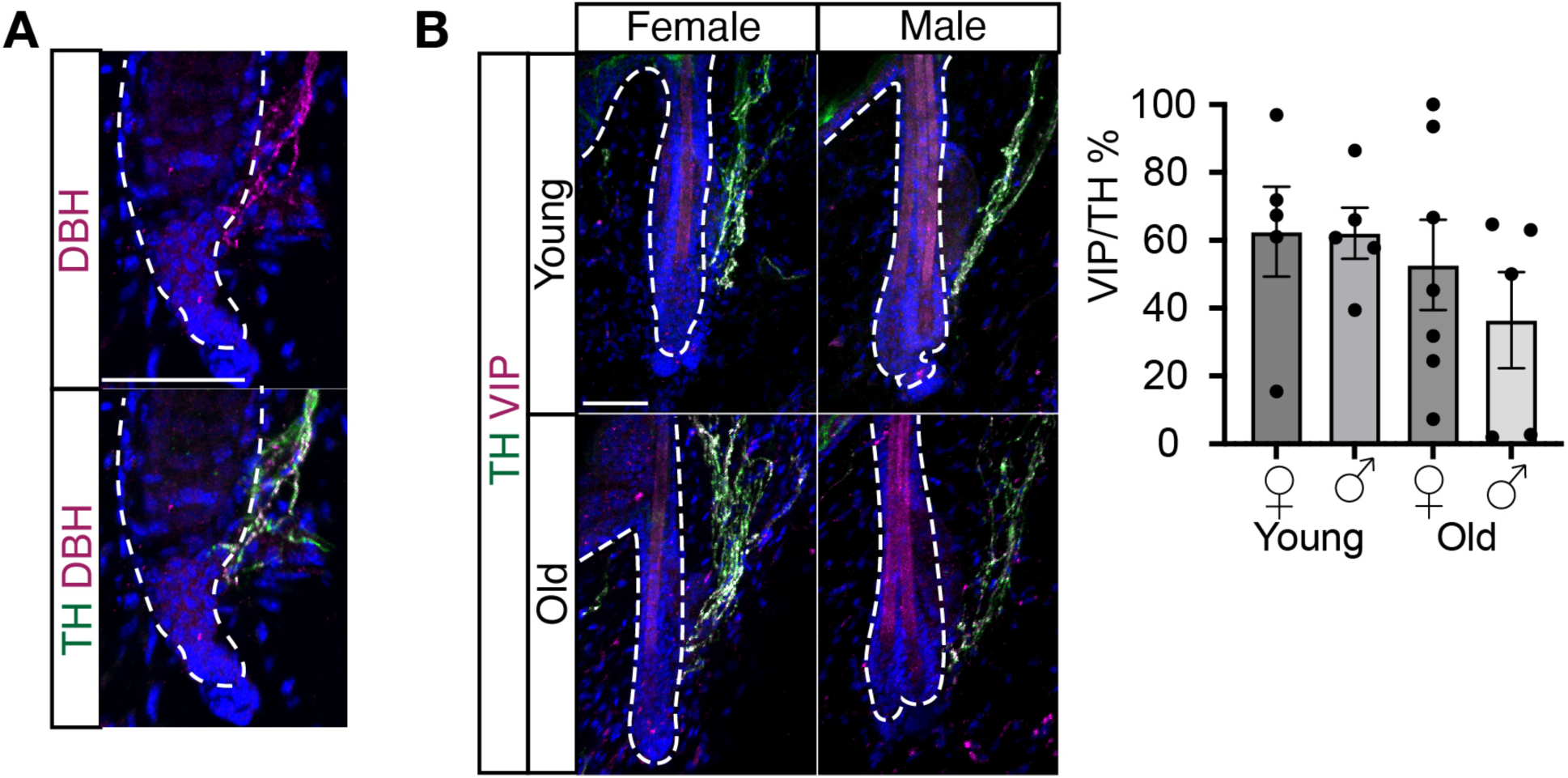
| Hair follicle-associated sympathetic adrenergic nerves maintain VIP expression during aging. (A) Representative images of tyrosine hydroxylase (TH; green) and dopamine β-hydroxylase (DBH; magenta) immunostaining demonstrating co-localization of TH and DBH in hair follicle-associated sympathetic nerve fibers. (B) Representative images of tyrosine hydroxylase (TH; green) and vasoactive intestinal peptide (VIP; magenta) immunostaining demonstrating co-localization of TH and VIP in hair follicle-associated sympathetic nerve fibers in young and old animals of both sexes. Quantification of VIP immunoreactivity in TH-positive sympathetic nerve fibers in young and old female and male mice. For (B), n = 5–7 mice per group, with 35–130 sympathetic nerve fibers analyzed per mouse. Each dot represents one mouse. Bars represent mean ± s.e.m. Statistical significance was determined using two-sided Student’s t-tests. Exact P values are shown. Scale bar, 50 µm.

**Supplementary Figure S3.**
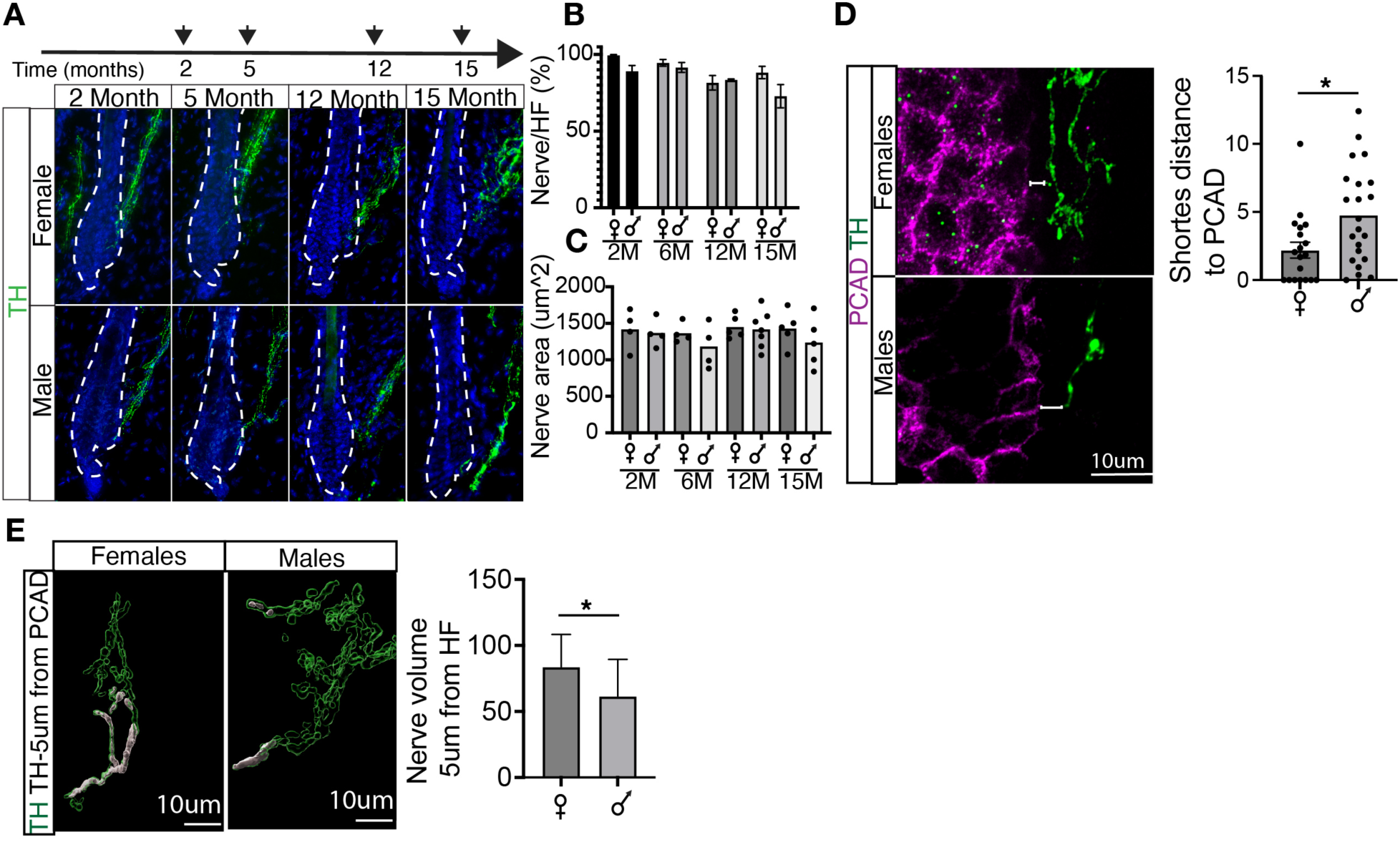
| Sex differences first emerge during neuroeffector remodeling rather than sympathetic nerve loss. (A) Representative images of tyrosine hydroxylase (TH; green) immunostaining in skin from 2-, 5-6, 12-, and 15-month-old female and male mice. (B) Quantification of the percentage of hair follicles associated with TH-positive sympathetic nerves (Nerve/HF) at 2, 5, 12, and 15 months of age. (C) Quantification of sympathetic nerve area at 2, 5, 12, and 15 months of age. (D) Representative images of TH-positive sympathetic nerves (green) and P-cadherin (PCAD; magenta) in 15-month-old female and male skin illustrating early alterations in neuroeffector organization. (E) Representative images of segmented TH-positive sympathetic nerves (green); white marks the nerve volume within 5 µm of the hair follicle. Right, quantification of sympathetic nerve volume within 5 µm of the hair follicle. For (B), n = 3–7 mice per group, with 20–90 hair follicles analyzed per mouse. Each dot represents one mouse. For (C), n = 4–7 mice per group, with 10 sympathetic nerves analyzed per mouse. Each dot represents one mouse. For (D) and (E), n = 4–5 mice per group, with 4–5 images analyzed per mouse. Each dot represents one image. Bars represent mean ± s.e.m. Statistical significance was determined using two-sided Mann–Whitney U tests for (B), (C), and (E) and a two-sided Student’s t-test for (D). Exact P values are shown. Scale bars: (A), 50 µm; (D-E) 10 µm.

**Supplementary Figure S4.**
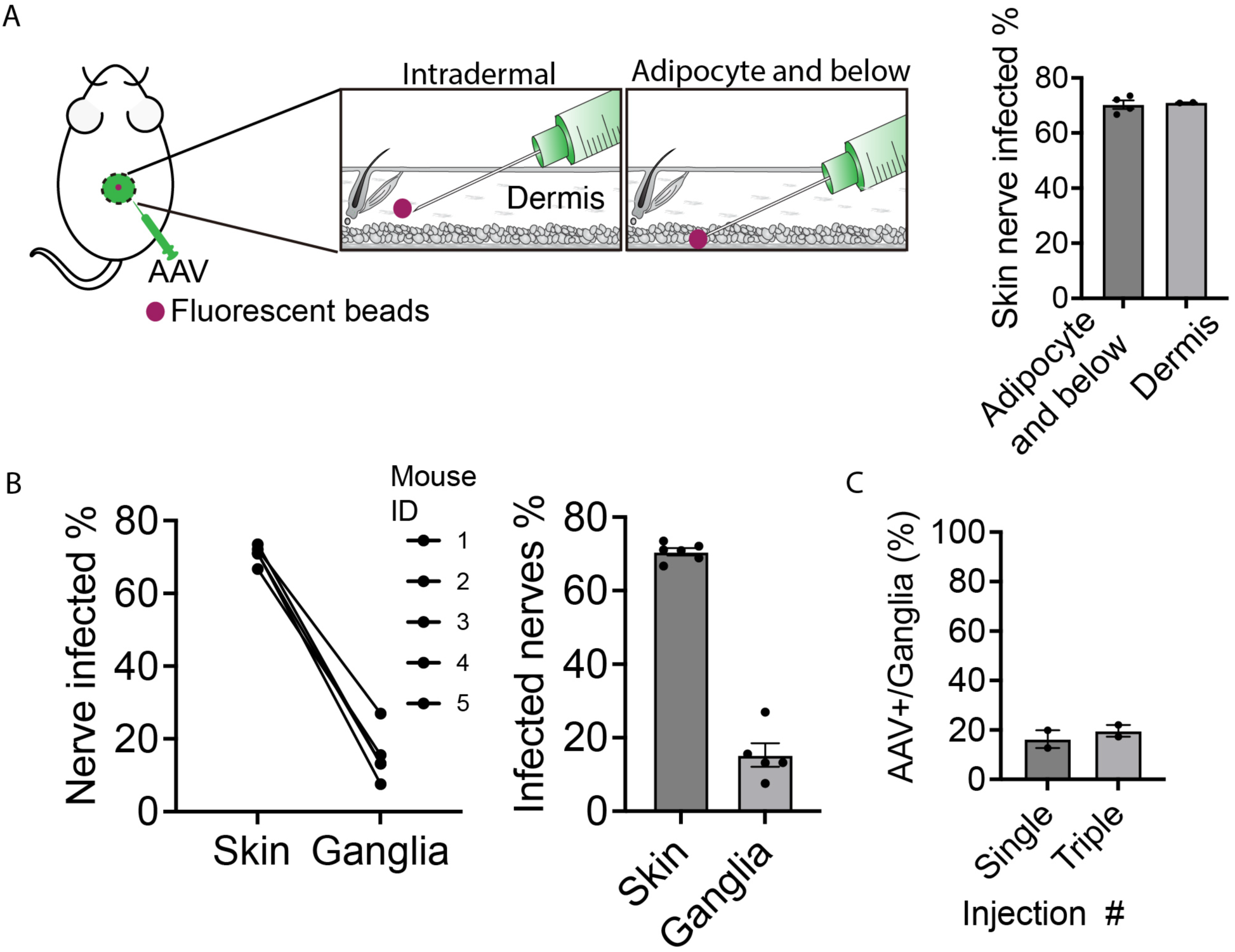
| Optimization of retrograde AAV labeling of skin-projecting sympathetic neurons. (A) Schematic illustrating intradermal and subdermal AAV-retro injection strategies. Fluorescent microspheres (magenta) were co-injected with AAV-retro to verify injection location. Right, quantification of the percentage of TH-positive sympathetic nerves labeled following intradermal or subdermal injection. Each dot represents one mouse, with 50–150 sympathetic nerves analyzed per mouse. (B) Reproducibility of retrograde AAV labeling. Left, paired comparison of the percentage of labeled sympathetic nerves in the skin and AAV-positive sympathetic neurons in thoracic sympathetic ganglia from individual mice following a single intradermal AAV-retro injection. Right, summary of labeling efficiencies in the skin and sympathetic ganglia. Each dot represents one mouse. (C) Comparison of retrograde labeling efficiency following one or three intradermal AAV-retro injections. Quantification shows the percentage of GFP-positive sympathetic neurons within thoracic sympathetic ganglia. Each dot represents one mouse, with 13–18 ganglion sections analyzed per mouse. Bars represent mean ± s.e.m. Statistical significance was determined using two-sided Mann– Whitney U tests.

**Supplementary Figure S5.**
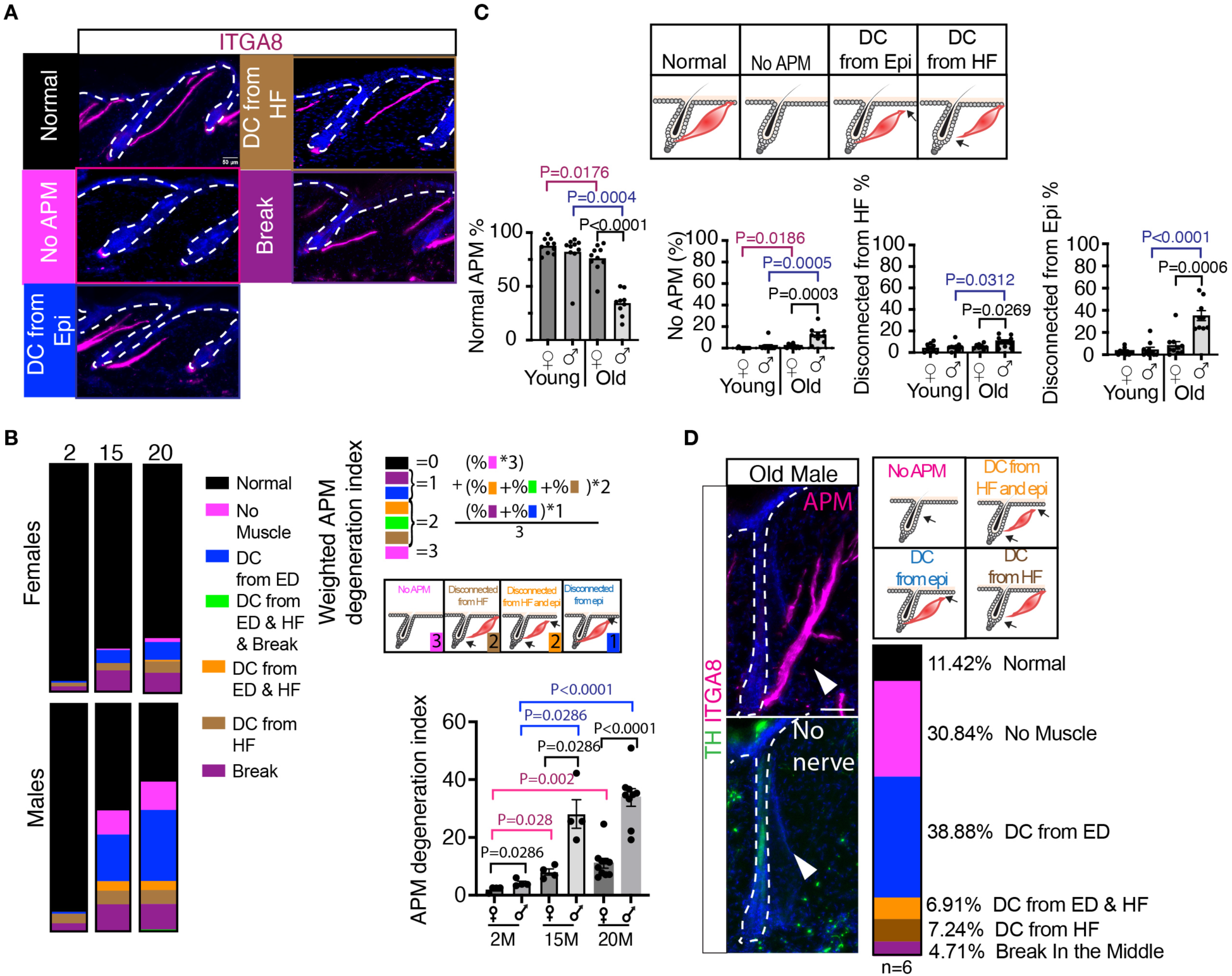
| Progressive degeneration of arrector pili muscles during aging. (A) Representative images of integrin α8 (ITGA8; magenta) immunostaining illustrating the major structural abnormalities of arrector pili muscles (APMs), including No APM, Disconnected from hair follicle (HF), Disconnected from epidermis (Epi), and Break. Dashed lines outline individual hair follicles. (B) Distribution of APM phenotypes in female and male mice at 2, 15, and 20 months of age. Right, schematic illustrating calculation of the weighted APM degeneration index. Bottom, quantification of the weighted APM degeneration index. (C) Schematic illustrating the major classes of APM degeneration. Right, quantification of the percentage of hair follicles exhibiting Normal APM, No APM, Disconnected from HF, and Disconnected from Epi in young and old female and male mice. (D) Representative images of tyrosine hydroxylase (TH; green) and ITGA8 (magenta) immunostaining in denervated hair follicles from aged male skin. Right, distribution of APM phenotypes among hair follicles lacking sympathetic innervation. For (B) and (C), n = 4–10 mice per group, with 40–80 hair follicles analyzed per mouse. Each dot represents one mouse. For (D), n = 6 aged male mice, with 10–25 denervated hair follicles analyzed per mouse. Bars represent mean ± s.e.m. Statistical significance was determined using two-sided Mann–Whitney U tests, except for Disconnected from HF in (C), which was analyzed using a two-sided Student’s t-test. Exact P values are shown. Pink brackets indicate age-related comparisons in females, blue brackets indicate age-related comparisons in males, and black brackets indicate comparisons between sexes. Scale bars: (A, D), 50 µm

**Supplementary Figure S6.**
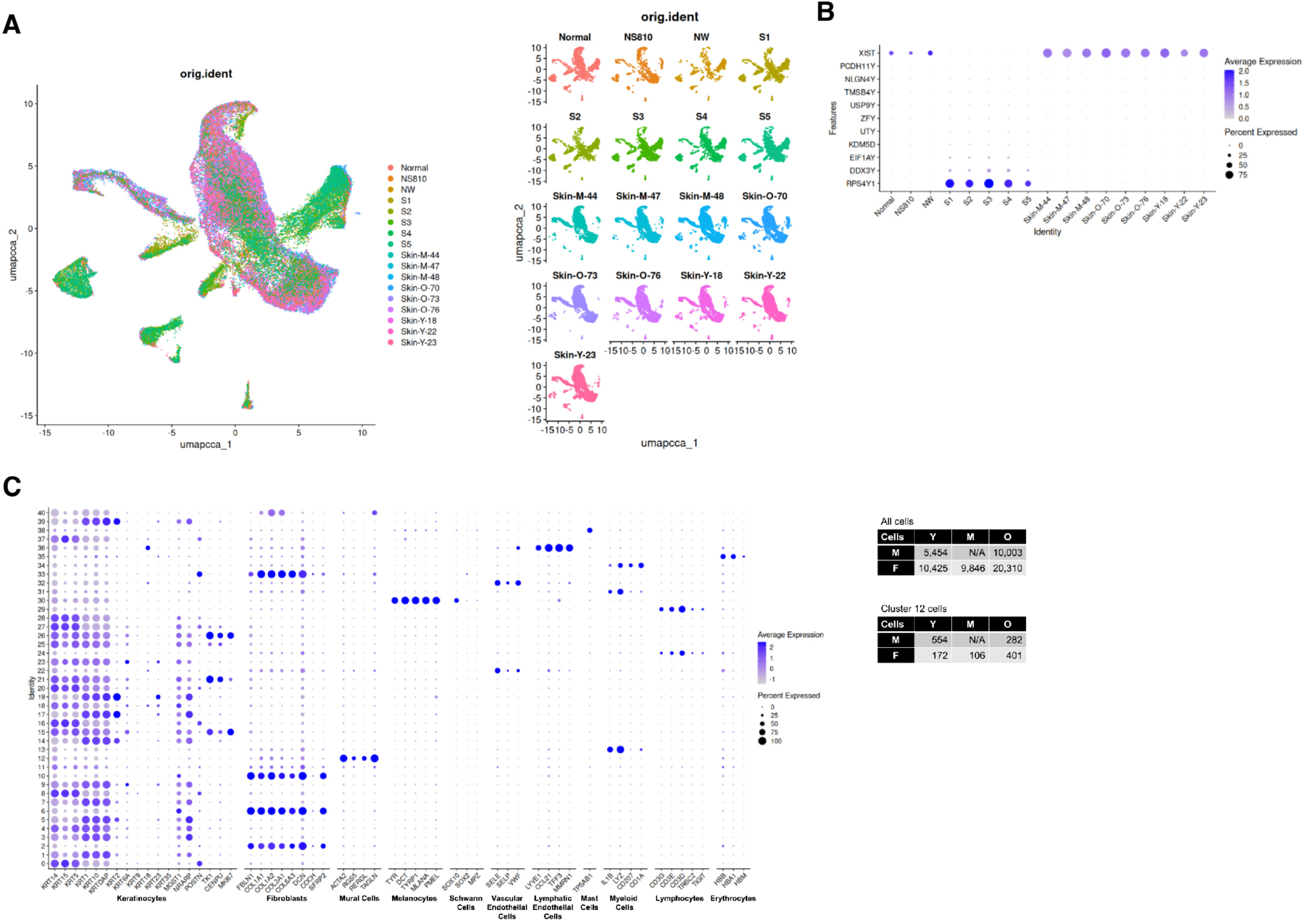
| Integration and annotation of human skin single-cell RNA-seq datasets. (A) Uniform manifold approximation and projection (UMAP) of the integrated human skin single-cell RNA-seq dataset colored by dataset of origin, demonstrating successful integration of samples from Hughes et al., Solé-Boldo et al., and Zou et al. Right, UMAPs showing the distribution of individual datasets after integration. (B) Dot plot showing expression of XIST and Y chromosome-associated genes used to confirm donor sex and verify the absence of testis-specific Y chromosome gene expression in skin samples. (C) Dot plot showing expression of canonical marker genes used to annotate the major cell populations in the integrated dataset. Tables summarize the numbers of young (Y), middle-aged (M), and aged (O) cells in the complete integrated dataset and in the smooth muscle cell cluster (cluster 12).

**Supplementary Figure S7.**
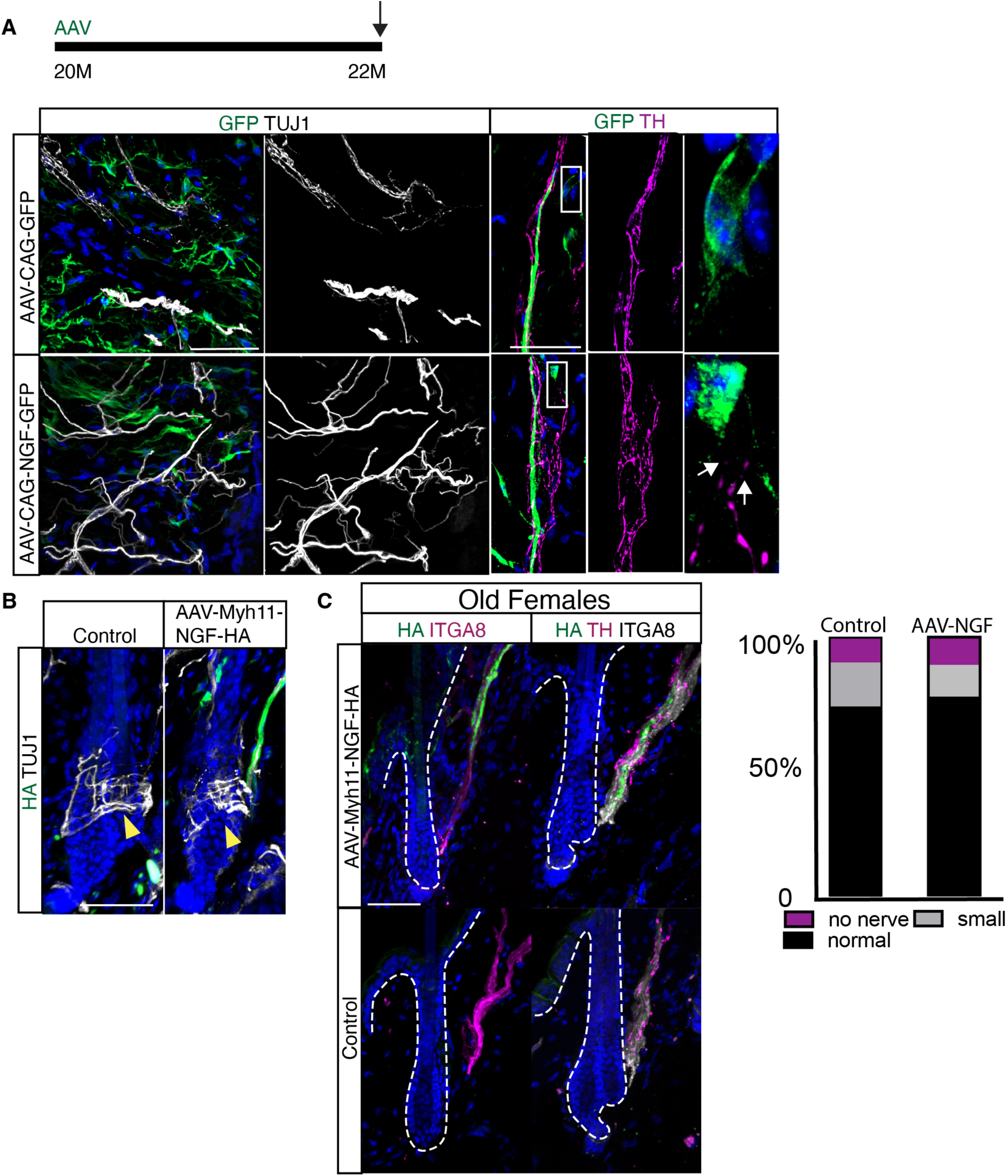
| Validation of broad and arrector pili muscle-specific NGF overexpression strategies. (A) Experimental design for broad NGF overexpression in aged mice using AAV8-CAG-NGF-T2A-GFP. Representative images of TUJ1-positive sensory nerve fibers (white) and GFP (green) in control (AAV-CAG-GFP) and NGF-treated (AAV-CAG-NGF-T2A-GFP) skin (left). Representative images of tyrosine hydroxylase (TH; magenta) and GFP (green) in control and NGF-treated skin (right). Insets show higher-magnification views of representative GFP-positive cells (B) Representative images of HA tag (green) and TUJ1 (white) immunostaining in control and AAV-Myh11-NGF-HA-treated skin demonstrating that arrector pili muscle-specific NGF overexpression does not disrupt hair follicle-associated sensory nerve terminals. Arrowheads indicate representative TUJ1-positive sensory nerve endings. (C) Representative images of HA (green) and ITGA8 (magenta) (left), and HA (green), TH (magenta), and ITGA8 (white) immunostaining (right) in control and AAV-Myh11-NGF-HA-treated aged female skin. Quantification shows no major difference in sympathetic nerve morphology following APM-specific NGF expression in aged females. Scale bars, 50 µm. For (A,B), n = 4 mice; for (C), n = 2 mice.

**Supplementary Figure S8.**
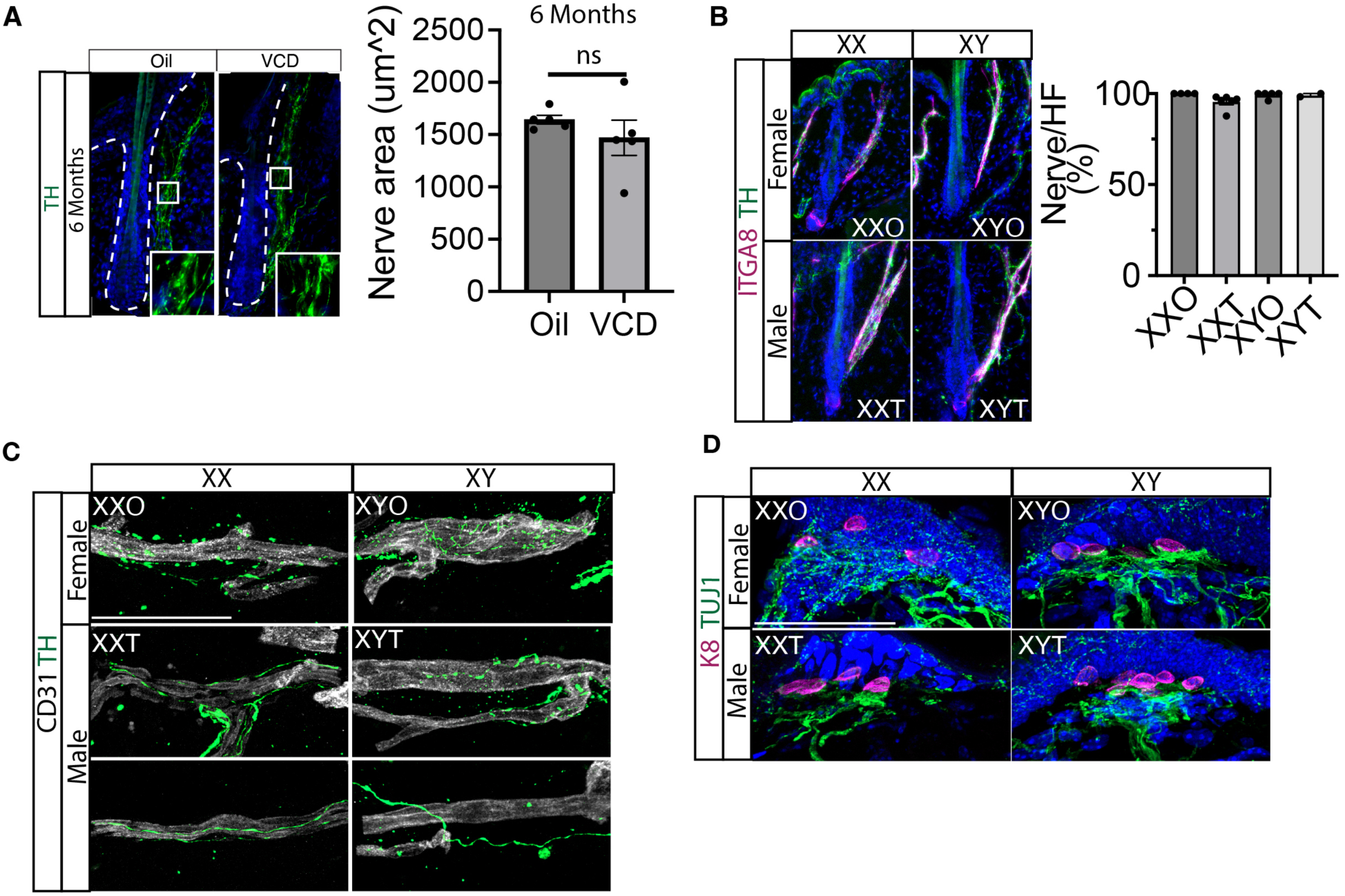
| Additional characterization of hormonal regulation of sympathetic aging. (A) Representative images of tyrosine hydroxylase (TH; green) immunostaining in skin from vehicle– and VCD-treated mice at 6 months of age. Insets show higher-magnification views of representative sympathetic nerve terminals. Right, quantification of sympathetic nerve area in 6-month-old mice following VCD treatment. (B) Representative images of tyrosine hydroxylase (TH; green) and ITGA8 (magenta) immunostaining in skin from four core genotype mice at 6-7 months. Quantification of the percentage of hair follicles with adjacent sympathetic nerves in a 6-7-month-old four-core genotype model. Each dot represents one mouse. XX0, XX animals with ovaries; XYO, XY animals with ovaries; XXT, XX animals with testis; XYT, XY animals with testis. (C) Representative images of TH (green) and CD31 (white) immunostaining in aged Four Core Genotypes mice illustrating sympathetic innervation of cutaneous blood vessels. (D) Representative images of TUJ1 (green) and keratin 8 (K8; magenta) immunostaining in aged Four Core Genotypes mice showing Merkel cell-associated sensory innervation. For (A), n =5 mice per group. Ten sympathetic nerves were analyzed per mouse. Each dot represents one mouse. Bars represent mean ± s.e.m. Statistical significance was determined using a two-sided Mann–Whitney U test. Scale bars, 50 µm. in C and D: XX females: XXO, XY females: XY0, XX males: XXT, XY males: XYT.

## References

1 Chavan, S. S., Pavlov, V. A. & Tracey, K. J. Mechanisms and Therapeutic Relevance of Neuro-immune Communication. Immunity 46, 927–942, doi:10.1016/j.immuni.2017.06.008 (2017).

2 Handler, A. & Ginty, D. D. The mechanosensory neurons of touch and their mechanisms of activation. Nat Rev Neurosci 22, 521–537, doi:10.1038/s41583-021-00489-x (2021).

3 Peng, J., Chen, H. & Zhang, B. Nerve-stem cell crosstalk in skin regeneration and diseases. Trends Mol Med 28, 583–595, doi:10.1016/j.molmed.2022.04.005 (2022).

4 Scott-Solomon, E. & Hsu, Y. C. Neurobiology, Stem Cell Biology, and Immunology: An Emerging Triad for Understanding Tissue Homeostasis and Repair. Annu Rev Cell Dev Biol 38, 419–446, doi:10.1146/annurev-cellbio-120320-032429 (2022).

5 Maryanovich, M. et al. Adrenergic nerve degeneration in bone marrow drives aging of the hematopoietic stem cell niche. Nat Med 24, 782–791, doi:10.1038/s41591-018-0030-x (2018).

6 Phillips, R. J., Hudson, C. N. & Powley, T. L. Sympathetic axonopathies and hyperinnervation in the small intestine smooth muscle of aged Fischer 344 rats. Auton Neurosci 179, 108–121, doi:10.1016/j.autneu.2013.09.002 (2013).

7 Masliukov, P. M., Emanuilov, A. I. & Budnik, A. F. Sympathetic innervation of the development, maturity, and aging of the gastrointestinal tract. Anat Rec (Hoboken) 306, 2249–2263, doi:10.1002/ar.25015 (2023).

8 Wagner, J. U. G. et al. Aging impairs the neurovascular interface in the heart. Science 381, 897–906, doi:10.1126/science.ade4961 (2023).

9 Carpenter, R. S. et al. Aging disrupts sympathetic innervation of the thymus. Cell Rep 45, 117126, doi:10.1016/j.celrep.2026.117126 (2026).

10 Chartier, S. R., Mitchell, S. A. T., Majuta, L. A. & Mantyh, P. W. The Changing Sensory and Sympathetic Innervation of the Young, Adult and Aging Mouse Femur. Neuroscience 387, 178–190, doi:10.1016/j.neuroscience.2018.01.047 (2018).

11 Hägg, S. & Jylhävä, J. Sex differences in biological aging with a focus on human studies. Elife 10, doi:10.7554/eLife.63425 (2021).

12 Sampathkumar, N. K. et al. Widespread sex dimorphism in aging and age-related diseases. Hum Genet 139, 333–356, doi:10.1007/s00439-019-02082-w (2020).

13 Hicks, C. W., Wang, D., Windham, B. G., Matsushita, K. & Selvin, E. Prevalence of peripheral neuropathy defined by monofilament insensitivity in middle-aged and older adults in two US cohorts. Sci Rep 11, 19159, doi:10.1038/s41598-021-98565-w (2021).

14 Lauria, G. et al. Intraepidermal nerve fiber density at the distal leg: a worldwide normative reference study. J Peripher Nerv Syst 15, 202–207, doi:10.1111/j.1529-8027.2010.00271.x (2010).

15 Davis, E. J., Lobach, I. & Dubal, D. B. Female XX sex chromosomes increase survival and extend lifespan in aging mice. Aging Cell 18, e12871, doi:10.1111/acel.12871 (2019).

16 Dart, A. M., Du, X. J. & Kingwell, B. A. Gender, sex hormones and autonomic nervous control of the cardiovascular system. Cardiovasc Res 53, 678–687, doi:10.1016/s0008-6363(01)00508-9 (2002).

17 Joyner, M. J., Barnes, J. N., Hart, E. C., Wallin, B. G. & Charkoudian, N. Neural control of the circulation: how sex and age differences interact in humans. Compr Physiol 5, 193–215, doi:10.1002/cphy.c140005 (2015).

18 Keir, D. A. et al. Influence of Sex and Age on Muscle Sympathetic Nerve Activity of Healthy Normotensive Adults. Hypertension 76, 997–1005, doi:10.1161/hypertensionaha.120.15208 (2020).

19 Klassen, S. A., Joyner, M. J. & Baker, S. E. The impact of ageing and sex on sympathetic neurocirculatory regulation. Semin Cell Dev Biol 116, 72–81, doi:10.1016/j.semcdb.2021.01.001 (2021).

20 Fujiwara, H. et al. The basement membrane of hair follicle stem cells is a muscle cell niche. Cell 144, 577–589, doi:10.1016/j.cell.2011.01.014 (2011).

21 Furlan, A. et al. Visceral motor neuron diversity delineates a cellular basis for nipple– and pilo-erection muscle control. Nat Neurosci 19, 1331–1340, doi:10.1038/nn.4376 (2016).

22 Honma, Y. et al. Artemin is a vascular-derived neurotropic factor for developing sympathetic neurons. Neuron 35, 267–282, doi:10.1016/s0896-6273(02)00774-2 (2002).

23 Damon, D. H., Teriele, J. A. & Marko, S. B. Vascular-derived artemin: a determinant of vascular sympathetic innervation? Am J Physiol Heart Circ Physiol 293, H266–273, doi:10.1152/ajpheart.00859.2006 (2007).

24 Crowley, C. et al. Mice lacking nerve growth factor display perinatal loss of sensory and sympathetic neurons yet develop basal forebrain cholinergic neurons. Cell 76, 1001–1011, doi:10.1016/0092-8674(94)90378-6 (1994).

25 Davis, B. M. et al. Overexpression of nerve growth factor in skin causes preferential increases among innervation to specific sensory targets. J Comp Neurol 387, 489–506, doi:10.1002/(sici)1096-9861(19971103)387:4<489::aid-cne2>3.0.co;2-z (1997).

26 Scott-Solomon, E., Boehm, E. & Kuruvilla, R. The sympathetic nervous system in development and disease. Nat Rev Neurosci 22, 685–702, doi:10.1038/s41583-021-00523-y (2021).

27 Diamond, J., Holmes, M. & Coughlin, M. Endogenous NGF and nerve impulses regulate the collateral sprouting of sensory axons in the skin of the adult rat. J Neurosci 12, 1454–1466, doi:10.1523/JNEUROSCI.12-04-01454.1992 (1992).

28 Glebova, N. O. & Ginty, D. D. Growth and survival signals controlling sympathetic nervous system development. Annu Rev Neurosci 28, 191–222, doi:10.1146/annurev.neuro.28.061604.135659 (2005).

29 Silver, J. & Miller, J. H. Regeneration beyond the glial scar. Nat Rev Neurosci 5, 146–156, doi:10.1038/nrn1326 (2004).

30 Fan, S. M. et al. External light activates hair follicle stem cells through eyes via an ipRGC-SCN-sympathetic neural pathway. Proc Natl Acad Sci U S A 115, E6880–e6889, doi:10.1073/pnas.1719548115 (2018).

31 Miranda, M. et al. Defining a Role for G-Protein Coupled Receptor/cAMP/CRE-Binding Protein Signaling in Hair Follicle Stem Cell Activation. J Invest Dermatol 142, 53–64.e53, doi:10.1016/j.jid.2021.05.031 (2022).

32 Shwartz, Y. et al. Cell Types Promoting Goosebumps Form a Niche to Regulate Hair Follicle Stem Cells. Cell 182, 578–593 e519, doi:10.1016/j.cell.2020.06.031 (2020).

33 Guo, M., Jiang, J., Zhang, A., Yu, W. & Huang, X. Cholesterol promotes hair growth through activating sympathetic nerves and enhancing the proliferation of hair follicle stem cells. Mol Med 31, 86, doi:10.1186/s10020-025-01139-z (2025).

34 Kumari, R. et al. Sympathetic NPY controls glucose homeostasis, cold tolerance, and cardiovascular functions in mice. Cell Rep 43, 113674, doi:10.1016/j.celrep.2024.113674 (2024).

35 Klassen, S. A. et al. Human sympathetic neuronal discharge and recruitment patterns regulate neuropeptide Y bioavailability. Am J Physiol Heart Circ Physiol 327, H1599–h1605, doi:10.1152/ajpheart.00639.2024 (2024).

36 Courtney, N. A. & Ford, C. P. The timing of dopamine– and noradrenaline-mediated transmission reflects underlying differences in the extent of spillover and pooling. J Neurosci 34, 7645–7656, doi:10.1523/JNEUROSCI.0166-14.2014 (2014).

37 Burnstock, G. Non-synaptic transmission at autonomic neuroeffector junctions. Neurochem Int 52, 14–25, doi:10.1016/j.neuint.2007.03.007 (2008).

38 Jänig, W. Functional Anatomy of the Peripheral Sympathetic and Parasympathetic Systems., 9–33 (Cambridge University Press, 2022).

39 Tervo, D. G. et al. A Designer AAV Variant Permits Efficient Retrograde Access to Projection Neurons. Neuron 92, 372–382, doi:10.1016/j.neuron.2016.09.021 (2016).

40 Schmidt, R. E. Age-related sympathetic ganglionic neuropathology: human pathology and animal models. Auton Neurosci 96, 63–72, doi:10.1016/s1566-0702(01)00372-1 (2002).

41 Ge, Y. et al. The aging skin microenvironment dictates stem cell behavior. Proc Natl Acad Sci U S A 117, 5339–5350, doi:10.1073/pnas.1901720117 (2020).

42 Hughes, T. K. et al. Second-Strand Synthesis-Based Massively Parallel scRNA-Seq Reveals Cellular States and Molecular Features of Human Inflammatory Skin Pathologies. Immunity 53, 878–894 e877, doi:10.1016/j.immuni.2020.09.015 (2020).

43 Sole-Boldo, L. et al. Single-cell transcriptomes of the human skin reveal age-related loss of fibroblast priming. Commun Biol 3, 188, doi:10.1038/s42003-020-0922-4 (2020).

44 Zou, Z. et al. A Single-Cell Transcriptomic Atlas of Human Skin Aging. Dev Cell 56, 383–397 e388, doi:10.1016/j.devcel.2020.11.002 (2021).

45 Aravamudan, B., Thompson, M., Pabelick, C. & Prakash, Y. S. Brain-derived neurotrophic factor induces proliferation of human airway smooth muscle cells. J Cell Mol Med 16, 812–823, doi:10.1111/j.1582-4934.2011.01356.x (2012).

46 Britt, R. D., Jr., et al. Smooth muscle brain-derived neurotrophic factor contributes to airway hyperreactivity in a mouse model of allergic asthma. FASEB J 33, 3024–3034, doi:10.1096/fj.201801002R (2019).

47 Prakash, Y. S., Iyanoye, A., Ay, B., Mantilla, C. B. & Pabelick, C. M. Neurotrophin effects on intracellular Ca2+ and force in airway smooth muscle. Am J Physiol Lung Cell Mol Physiol 291, L447–456, doi:10.1152/ajplung.00501.2005 (2006).

48 Roos, B. B. et al. Neurotrophin Regulation and Signaling in Airway Smooth Muscle. Adv Exp Med Biol 1304, 109–121, doi:10.1007/978-3-030-68748-9_7 (2021).

49 Vohra, P. K. et al. TRPC3 regulates release of brain-derived neurotrophic factor from human airway smooth muscle. Biochim Biophys Acta 1833, 2953–2960, doi:10.1016/j.bbamcr.2013.07.019 (2013).

50 Freeman, M. R. et al. Brain-derived neurotrophic factor and airway fibrosis in asthma. Am J Physiol Lung Cell Mol Physiol 313, L360–L370, doi:10.1152/ajplung.00580.2016 (2017).

51 Monica Brauer, M. & Smith, P. G. Estrogen and female reproductive tract innervation: cellular and molecular mechanisms of autonomic neuroplasticity. Auton Neurosci 187, 1–17, doi:10.1016/j.autneu.2014.11.009 (2015).

52 Wyss, J. M. & Carlson, S. H. Effects of hormone replacement therapy on the sympathetic nervous system and blood pressure. Curr Hypertens Rep 5, 241–246, doi:10.1007/s11906-003-0027-8 (2003).

53 Brooks, H. L., Pollow, D. P. & Hoyer, P. B. The VCD Mouse Model of Menopause and Perimenopause for the Study of Sex Differences in Cardiovascular Disease and the Metabolic Syndrome. Physiology (Bethesda) 31, 250–257, doi:10.1152/physiol.00057.2014 (2016).

54 Arnold, A. P. Four Core Genotypes and XY* mouse models: Update on impact on SABV research. Neurosci Biobehav Rev 119, 1–8, doi:10.1016/j.neubiorev.2020.09.021 (2020).

55 Arnold, A. P. & Chen, X. What does the “four core genotypes” mouse model tell us about sex differences in the brain and other tissues? Front Neuroendocrinol 30, 1–9, doi:10.1016/j.yfrne.2008.11.001 (2009).

56 Corre, C. et al. Separate effects of sex hormones and sex chromosomes on brain structure and function revealed by high-resolution magnetic resonance imaging and spatial navigation assessment of the Four Core Genotype mouse model. Brain Struct Funct 221, 997–1016, doi:10.1007/s00429-014-0952-0 (2016).

57 Kenney, M. J. Animal aging and regulation of sympathetic nerve discharge. J Appl Physiol (1985) 109, 951–958, doi:10.1152/japplphysiol.00506.2010 (2010).

58 Hart, E. C. et al. Sex and ageing differences in resting arterial pressure regulation: the role of the β-adrenergic receptors. J Physiol 589, 5285–5297, doi:10.1113/jphysiol.2011.212753 (2011).

59 Hinojosa-Laborde, C., Chapa, I., Lange, D. & Haywood, J. R. Gender differences in sympathetic nervous system regulation. Clin Exp Pharmacol Physiol 26, 122–126, doi:10.1046/j.1440-1681.1999.02995.x (1999).

60 Reckelhoff, J. F. Sex Differences in Regulation of Blood Pressure. Adv Exp Med Biol 1065, 139–151, doi:10.1007/978-3-319-77932-4_9 (2018).

61 Blake, M. R. et al. Loss of chondroitin sulfate proteoglycan sulfation allows delayed sympathetic reinnervation after cardiac ischemia-reperfusion. Physiol Rep 11, e15702, doi:10.14814/phy2.15702 (2023).

62 Lorentz, C. U. et al. Sympathetic denervation of peri-infarct myocardium requires the p75 neurotrophin receptor. Exp Neurol 249, 111–119, doi:10.1016/j.expneurol.2013.08.015 (2013).

63 Hsu, Y. C. & Fuchs, E. Building and Maintaining the Skin. Cold Spring Harb Perspect Biol 14, doi:10.1101/cshperspect.a040840 (2022).

64 Li, K. N. & Tumbar, T. Hair follicle stem cells as a skin-organizing signaling center during adult homeostasis. EMBO J 40, e107135, doi:10.15252/embj.2020107135 (2021).

65 Zhang, B. & Chen, T. Local and systemic mechanisms that control the hair follicle stem cell niche. Nat Rev Mol Cell Biol 25, 87–100, doi:10.1038/s41580-023-00662-3 (2024).

66 Shin, S. H., Lee, Y. H., Rho, N. K. & Park, K. Y. Skin aging from mechanisms to interventions: focusing on dermal aging. Front Physiol 14, 1195272, doi:10.3389/fphys.2023.1195272 (2023).

67 Tobin, D. J. Introduction to skin aging. J Tissue Viability 26, 37–46, doi:10.1016/j.jtv.2016.03.002 (2017).

68 Jang, H., Jo, Y., Lee, J. H. & Choi, S. Aging of hair follicle stem cells and their niches. BMB Rep 56, 2–9, doi:10.5483/BMBRep.2022-0183 (2023).

69 Giangreco, A., Qin, M., Pintar, J. E. & Watt, F. M. Epidermal stem cells are retained in vivo throughout skin aging. Aging Cell 7, 250–259, doi:10.1111/j.1474-9726.2008.00372.x (2008).

70 Peters, E. M. et al. Nerve growth factor and its precursor differentially regulate hair cycle progression in mice. J Histochem Cytochem 54, 275–288, doi:10.1369/jhc.4A6585.2005 (2006).

71 Botchkarev, V. A., Botchkareva, N. V., Peters, E. M. & Paus, R. Epithelial growth control by neurotrophins: leads and lessons from the hair follicle. Prog Brain Res 146, 493–513, doi:10.1016/S0079-6123(03)46031-7 (2004).

72 Habecker, B. A. et al. Molecular and cellular neurocardiology in heart disease. J Physiol 603, 1689–1728, doi:10.1113/JP284739 (2025).

73 Pellegrino, M. J. & Habecker, B. A. STAT3 integrates cytokine and neurotrophin signals to promote sympathetic axon regeneration. Mol Cell Neurosci 56, 272–282, doi:10.1016/j.mcn.2013.06.005 (2013).

74 Xie, X. et al. Hyperactivation of sympathetic nerves fuels basophil infiltration in atopic dermatitis. Immunity 59, 598–617.e511, doi:10.1016/j.immuni.2026.01.010 (2026).

75 Yu, S. et al. Sympathetic nerve dysfunction exacerbates skin inflammation in atopic dermatitis. J Allergy Clin Immunol 157, 677–692, doi:10.1016/j.jaci.2025.12.994 (2026).

76 Tian, J. et al. A sympathetic-eosinophil axis orchestrates psychological stress to exacerbate skin inflammation. Science 391, 1269–1277, doi:10.1126/science.adv5974 (2026).

77 Gao, X., Zhang, J. & Tamplin, O. J. The aging hematopoietic stem cell niche: a mini review. Front Hematol 4, doi:10.3389/frhem.2025.1525132 (2025).

78 Ho, Y. H. & Mendez-Ferrer, S. Microenvironmental contributions to hematopoietic stem cell aging. Haematologica 105, 38–46, doi:10.3324/haematol.2018.211334 (2020).

79 Xiang, Y. et al. Early menarche and childbirth accelerate aging-related outcomes and age-related diseases: Evidence for antagonistic pleiotropy in humans. Elife 13, doi:10.7554/eLife.102447 (2025).

80 Hamlat, E. J., Prather, A. A., Horvath, S., Belsky, J. & Epel, E. S. Early life adversity, pubertal timing, and epigenetic age acceleration in adulthood. Dev Psychobiol 63, 890–902, doi:10.1002/dev.22085 (2021).

81 Maddock, J. et al. Childhood growth and development and DNA methylation age in mid-life. Clin Epigenetics 13, 155, doi:10.1186/s13148-021-01138-x (2021).

82 Goering, M., Barker-Kamps, M., Patki, A., Tiwari, H. K. & Mrug, S. Pubertal timing as a predictor of epigenetic aging and mortality risk in young adulthood. Dev Psychol 61, 912–927, doi:10.1037/dev0001903 (2025).

83 Moses, E. et al. An antagonistically pleiotropic gene regulates vertebrate growth, maturity, and lifespan. Nat Commun 17, doi:10.1038/s41467-026-72381-0 (2026).

84 Panten, J. et al. Four Core Genotypes mice harbour a 3.2MB X-Y translocation that perturbs Tlr7 dosage. Nat Commun 15, 8814, doi:10.1038/s41467-024-52640-8 (2024).

85 Tam, H. T. et al. Hyperinnervation inhibits organ-level regeneration in mammalian skin. Cell 189, 3270–3286.e3217, doi:10.1016/j.cell.2026.02.027 (2026).

